# Interaction between climatic variation and pathogen diversity shape endemic disease dynamics in the agricultural settings

**DOI:** 10.1101/2024.09.27.615185

**Authors:** Rishi Bhandari, Amanpreet Kaur, Ivory Russell, Ogonnaya Michael Romanus, Destiny Brokaw, Anthony P Keinath, Zachary Snipes, Philip Rollins, Nettie Baugher, Inga Meadows, Ty Nicholas Torrance, Bhabesh Dutta, Edward Sikora, Roberto Molinari, Samuel Soubeyrand, Neha Potnis

## Abstract

Recurring outbreaks caused by endemic pathogens continue to pose problems in managing plant, wildlife, livestock and human health. Understanding how these outbreaks unfold and what drives the variability in disease epidemics across space and time is less understood, especially in the agricultural settings. In this study, we investigated the contribution of pathogen genetic diversity, climatic variation and their interaction towards disease dynamics, with an integrative approach grounded on multitype, high resolution sequencing data and analysis techniques. This investigation was carried out for bacterial spot disease epidemic by surveying tomato fields for bacterial pathogen (*Xanthomonas perforans*) across southeastern US over a span of three years. The strength of epidemic severity varied across space and time in the agricultural fields. Disease severity was positively associated with strain diversity, and was linked to environmental fluctuations, specifically, large variation and extreme changes in certain climatic factors. Strain-resolved metagenomics approach revealed that co-existence of multiple pathogen lineages was common in the individual fields, although accompanied by differential lineage dynamics. The co-occurring lineages displayed environmentally dependent fitness contributions. By tracing allelic frequencies in pathogen populations across temporal scales, we find evidence for asynchronous allele cycling across seasons, hinting at the presence of adaptive single-nucleotide polymorphisms (SNPs) being polymorphic in space in response to seasonality. Despite this pathogen heterogeneity, we identified positively selected loci under parallel evolution, which may explain the nature of selection pressures experienced by the pathogen. While single pathogen lineage is assumed to dominate the host in the agricultural settings, our findings challenge this notion by demonstrating genetic diversity in the pathogen population observed within a single field and linking it to the disease dynamics. Our results explain the role of pathogen genetic diversity, climate-dependent compositional dynamics, and differential fitness contributions in dictating the variability of disease epidemics in the agricultural settings. Such findings will be invaluable for building predictive models in disease epidemiology. Our high-resolution combinatorial approach exploiting high-resolution sequence data, metadata types and analysis tools, is general enough to finely investigate disease epidemics at large scales in diverse case-studies concerning plant, animal and human health.

## Introduction

Predicting the disease epidemics in the face of climate change is a major challenge in epidemiological studies of diseases impacting human, wildlife and plant health (1–4). Disease epidemics result from complex host-pathogen-environment interactions where ecological and evolutionary processes shape host and pathogen populations and environment can directly impact pathogen fitness or indirectly impact host susceptibility (5–9). These complexities involving interdependent interactions pose a challenge in dissecting the contribution of each component towards variability in disease epidemics. In addition, while controlled experiments have provided insights into specific environmental parameters in shaping disease outbreaks, we have little experimental evidence on how frequent shifts in these climatic factors can impact disease dynamics and whether climate change has already led to a surge of plant disease outbreaks over the past century (10–15).

Among different factors contributing to variable incidences and trajectories of disease epidemics, pathogen diversity plays a major role in this equation. While molecular markers and next-generation sequencing approaches have accelerated characterization of pathogen diversity, the diversity studies have been focused on informing pathogen population structure and evolution of the pathogen (16–19). Integral role of population genetics into epidemiology and disease management has been much appreciated with the rapid next-generation sequencing efforts (20,21). Time-series isolate-genome sequencing has transformed our views on population genetics going beyond conventional diagnostic tools informing races, pathotypes, or species (17), although isolation-based approaches present a bias of choosing the most dominant pathogen genotype for sequencing and thus, may suppose pathogen homogeneity in a diseased sample, leaving out the details of low abundant pathogen variants that may be co-occurring. Such approaches may miss the “true genetic diversity” existing within single field, or a single organ, or a single lesion. In case of agricultural systems, majority of the focus has been on the dominant pathogen genotypes for designing management strategies, specifically breeding efforts, and characterization of pathogen spatial genetic structure studies at the micro-geographical scale are limited. Two recent studies have indicated presence of polyclonal infections in a single disease lesion (22), and epidemics with multiple co-existing pathogen genotypes (23) in a single field. These findings underscore the need for characterizing spatio-temporal pathogen population structure and the strain dynamics at the micro-geographical scale as disease epidemics unfold (24).

While much of the work in understanding the disease dynamics and pathogen diversity across space and time has been conducted in the natural settings with wild plant pathogens (25,26), studies linking pathogen diversity and dynamics to explaining disease epidemics in the agricultural settings are limited (27–29). Many factors associated with modern agricultural practices such as global trade, use of chemicals, or intensified monoculture system can predispose the system susceptible to the disease epidemics. For example, in the case of seed-borne pathogens, the global trade facilitating the continuous input of different pathogen genotypes on seeds or plant materials circulated worldwide has been a driving force in the pathogen diversity observed under field conditions (30). Thus, the initial strain composition of pathogen population entering the agricultural fields can be variable, depending on the source of the inoculum on seeds, or transplants, or surviving population from the previous epidemic season in the fields (31–33). Upon establishment of these initial pathogen populations, variation in the pathogen dynamics across different fields depends on many factors. These include differential fitness levels of pathogen genotypes on the host considering both virulence and aggressiveness, interactions between pathogen and host genotype (host specificity), strain competition, climate sensitivity of individual pathogen genotypes, and other processes such as dispersal that may introduce new pathogen genotypes on to the plant during the course of the season (34–39). In addition, local conditions such as neighboring plants, may affect the strain assembly processes (40). Grower’s practices such as use of copper sprays, or other chemical pesticides can also impact strain composition, facilitating dominance of chemically resistant population (41–43), while managing fitness trade- offs with virulence. The field landscape can influence pathogen population structure (44). In addition to the ecological dynamics, pathogen populations also undergo diversification through a variety of genetic processes such as mutation or recombination, although studies characterizing these genetic processes within a span of single or consecutive growing seasons are limited (39,45). Higher genetic diversity in the pathogen population has been hypothesized to reflect higher evolutionary potential (46), to facilitate genetic exchange (22) and was shown to contribute to epidemic growth (24) and in some cases, higher genetic diversity was shown to affect transmission dynamics (26).

The pathosystem of bacterial leaf spot (BLS) on tomatoes caused by *Xanthomonas perforans (Xp)* (also referred to as *Xanthomonas euvesicatoria pv. perforans*) is an example of an endemic disease where no known host resistance genes have been deployed in commercial tomato cultivars and disease epidemics continue to be observed across all the tomato producing regions (47). Genome sequencing of time-series collection of field isolates have informed that extensive shifts in the pathogen population, including species displacement and extensive diversification have been observed in the past decades (23,31,33,47–50). Tomato production system that introduces multiple pathogen genotypes into the fields via transplants has been shown to shape pathogen population structure in the field (31). In addition, inoculum from the previous growing season may also be additional source of introduction, depending on the cultural practices in the field. However, assuming variable probabilities of introductions of pathogen genotypes into the field, the following questions still remains unanswered: Does a single pathogen genotype dominate the agricultural fields as it is assumed for implementing management strategies? What factors explain variation in disease dynamics observed across the fields? How does pathogen respond to abrupt changes in the climatic patterns? We addressed these questions in the present study by specifically focusing on the contribution of climatic variation, pathogen diversity and strain dynamics in explaining different bacterial spot epidemic trajectories in the agricultural field outbreaks. Using strain- resolved metagenomics to characterize pathogen diversity within a single field (23), we surveyed tomato fields in the southeastern US over the course of growing season in three consecutive years and found that multiple (up to 5) pathogen genotypes co-exist and their abundance varies across the fields. While at least three different pathogen genotypes seemed to dominate the southeastern US, their differential abundance across fields and across seasons was explained by differential impact of abrupt climatic changes on the fitness of the three genotypes. The findings from this work on strain dynamics and fitness will be useful in diversity-informed epidemiological predictions, especially under changing climate.

## Results

### Varying levels of BLS disease severity exist within the Southeastern US

A total of 66 tomato phyllosphere samples were gathered over three years (2020–2022) from Alabama, Georgia, North Carolina, and South Carolina (Fig 1A; S1A Table). The sampling method was determined based on the field size and the amount of disease present (Fig 1B). Metagenomic DNA from the leaf samples was subjected to shotgun metagenomics to obtain a high-resolution pathogen population structure in the southeastern US. Apart from five samples showing no BLS symptoms, the sampled fields showed variable levels of disease severity (S1B Table) across consistently sampled individual fields (S1 Fig), across the season and within and across states (Figs 1C and S2A). We observed that average disease severity in samples from Alabama was lower during mid-season compared to other states. But the average disease severity during the end season was higher in Alabama compared to other states (Figs 1C and S2A). However, disease severity was not consistently higher as the season progressed, as seen with samples from Georgia and South Carolina (Figs 1C and S2A). We used the SNV profiling method, strainEST, to detect pathogen lineages at finer resolution including those with low abundance levels. For this, we screened metagenomic reads for the presence of BLS pathogenic species, *Xp*, *Xeu (Xanthmonas euvesicatoria pv. euvesicatoria)*, and *Xeu-*related pathovars (*Xeu* sister clades). We observed the absence of *Xeu* in sampled tomato fields suggesting that *Xp* is the dominant pathogen of BLS in tomatoes in the southeastern United States. Using correlation analysis as an exploratory tool, we found that the absolute abundance of *Xp* (Figs 1D and S2B) positively correlated with disease severity (mid-season: R^2^ = 0.381, *p* < 0.001, end-season: R^2^ = 0.283, *p* < 0.01). A similar observation was made with the relative abundance of *Xp* (S2C Fig) (mid-season: R^2^ = 0.508, *p* < 0.001, end-season: R^2^ = 0.378, *p* < 0.001). The pathogen population was significantly associated with disease severity during the early season compared to the late season based on R^2^ value. Although small-scale farms showed slightly higher disease severity, this difference compared to commercial-scale farms was not statistically significant (*p* = 0.45, see S3A Fig).

**Fig 1.**
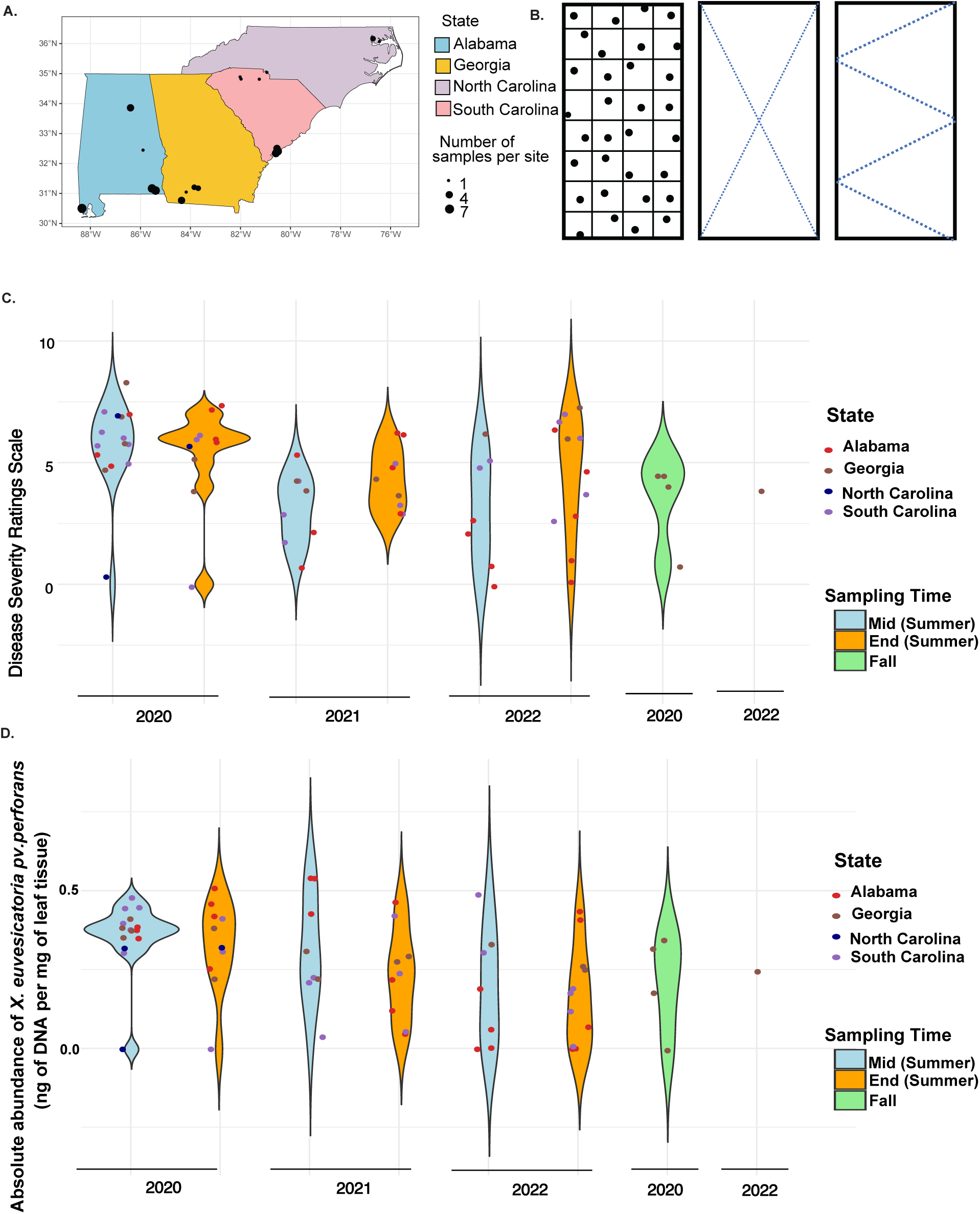
*Xanthomonas euvesicatoria* pv. *perforans* (*Xp*) continues to be a significant pathogen in the tomato fields across *the* southeastern US, although with variable disease severity in the region. **(A)** Map of the study area from Alabama, North Carolina, South Carolina, and Georgia from Southeast United States. The dots in the figures represent the sampled farm, and the size of the dot represents the number of samples collected during the years 2020, 2021, and 2022. The samples were collected from two different growing seasons: summer (with mid and end-season sampling points) and fall season. **(B)** Sampling methods varied by field size. In small farms, stratified random sampling was used to collect representative leaflets (left). In large fields, samples were taken along an X (mid) or W (right) pattern. Split-violin plots showing **(C)** disease severity ratings for all collected samples estimated using the Horsfall-Barratt scale, which ranges from 1 to 12, with the scale of 1 being no disease and 12 being 100% defoliation (51) and **(D)** the absolute abundance of *Xp* collected from different states during 2020, 2021, and 2022. The absolute abundance for each sample was estimated by multiplying the amount of DNA per ng of the sample with relative abundance of *Xp* present in it.

### *Xanthomonas* abundance reveals a heterogeneous spatial distribution of pathogen lineages, and the increased pathogen diversity was associated with higher disease severity

Given our observation of varying levels of epidemics, we hypothesized that pathogen heterogeneity is responsible for varying disease dynamics. Thus, we assessed the heterogeneity in the pathogen population by studying the extent of diversity of *Xp* in individual fields. Our approach involved mapping the metagenome reads to the SNP matrices of known pathogen lineages or sequence clusters (SC) of *Xp* and calculating relative abundance for each SC. To understand the current global population structure of *Xp*, we leveraged the availability of extensive genome resources for *Xp* collected worldwide over the last three decades (473 genomes as of March 2023). Based on the phylogeny of these publicly available global *Xp* collections, determined by core genome single nucleotide polymorphisms (SNPs), eight distinct lineages were identified within *Xp* strains referred to hereafter as sequence clusters (SC) or lineages (Fig 2A). This SNP-level resolution into different lineages was a baseline for our analysis to identify diversity and co-occurrence of pathogen lineages across field metagenomes. Surprisingly, we identified the presence of eight lineages in the field samples. Distantly related SC3 and 4 (SNP distance of 10,825 SNPs) co-existed in majority of the fields (Fig 2B). Three lineages, SC3, SC4, and SC6 were in higher abundance in these fields, although their abundance varied from field to field, and their dynamics varied across neighboring fields (Figs 2C and S4). Notably, SC1, SC7, and SC8 were in lower abundance, appearing in only eight samples (Figs 2C and S4). The co-occurrence of different *Xp* lineages were observed in the samples, as noted in our previous limited metagenome survey (23). Fields had a minimum of 1 to a maximum of 5 lineages present at a given sampling time, and this diversity of co-occurring lineages was different across different years, and samples from neighboring states (Figs 2C and S4). Next, we investigated whether multiple co-occurring lineages of *Xp* within individual fields influenced disease severity. Fields with more than two co-occurring SCs displayed higher average nucleotide diversity, and showed significantly higher disease severity compared to those with less than 2 SCs (S5 Fig). Correlation analysis between the Shannon diversity index of the pathogen population and disease severity revealed a positive association (R^2^ = 0.52, *p* < 0.001) (S6 Fig). This suggests that *Xp* population diversity is positively associated with disease severity. Changes in strain composition were not only evident across different fields but also within individual fields throughout the growing season, spanning three years. These dynamics in strain composition were marked by strain fluctuations, where the abundance of lineages shifted from one season to the other, strain invasion where a new lineage likely infected the plants during the growing season, and differential strain dynamics (Figs 2C and S4). Overall, the diversity of *Xp* found globally is circulating in Southeastern US tomato fields, and strain diversity and composition is significantly associated with disease severity.

**Fig 2.**
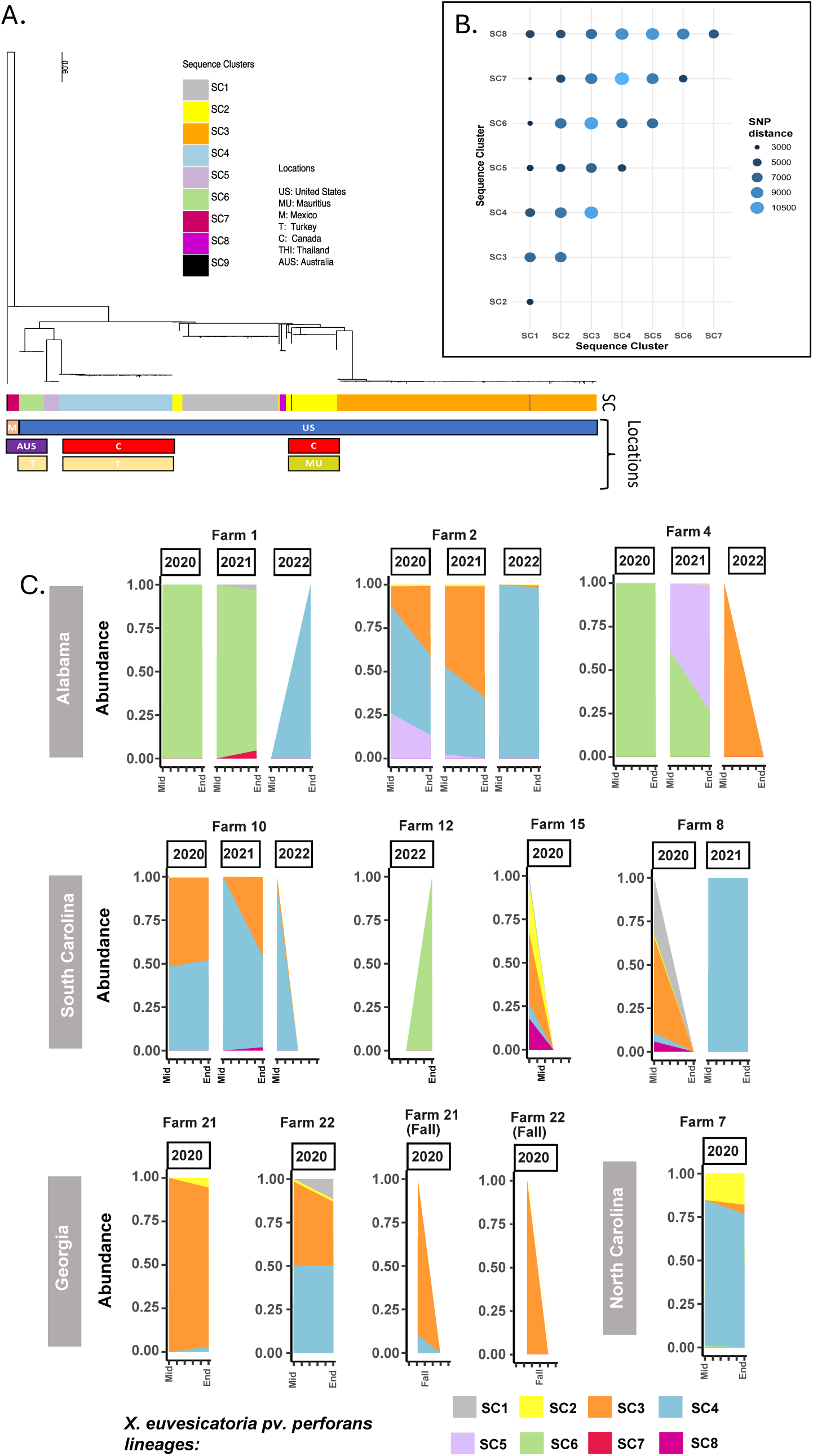
Co-occurrence of multiple lineages of the BLS pathogen observed in the southeastern United States, which leads to increased BLS disease severity. **(A)** Midpoint-rooted maximum-likelihood phylogeny based on a concatenated alignment of core genome SNPs from 473 *Xp* strains. The tips are color-coded according to the lineages identified in the first level of the HierBAPS hierarchy. SC9 is a single-strain lineage exclusively found in Nigeria. **(B)** Dot plot shows the SNP differences across different SCs, where size and colour of each point refer to the number of SNPs differences present while comparing one SC (on x-axis) comparison to another SC (on y-axis). **(C)** Stacked bar plot depicting the co-occurrence of multiple *Xp* lineages, spatial and temporal variations, the introduction of new lineages, turnover, and dominance shifts in individual fields across various states during the mid and end of summer seasons and fall seasons for the years 2020, 2021, and 2022. Representative fields are shown in this figure. **S4 Fig** is showing data for all fields is included in the supplementary file.

### Climatic shifts and extremes explain variation in BLS epidemics across the Southeastern United States

Given the findings of variable disease dynamics across neighboring states, we assessed the contribution of climatic factors and their spatial and temporal variations as drivers of disease epidemics. We used regression models to analyze how disease severity and pathogen abundance (response variables) are related to predictor variables such as climatic factors, sampling time, year, and farm scale (treating the latter three as categorical variables). While the data on disease severity, pathogen abundance, and pathogen diversity were collected for specific timepoints during the growing season, the datasets obtained from NASA-POWER for weather related variables (S2 Table) were at a much finer spatio-temporal resolution with daily record (0.5 x 0.625 latitude-longitude grid cell resolution). In addition to considering routine mean or median values of climatic variables for analyses, we also considered characteristics of climatic predictors using indexes that capture the spread (standard deviation), asymmetries, and tail-heaviness (skewness and kurtosis). This approach accounts for within-season climatic variations, considering interseasonal weather events, that may present highly conducive conditions for bacterial diseases. Considering the sample size, this approach, however, considerably increased the number of predictors so, to achieve reasonable statistical conclusions, for all regressions we ran a preliminary predictor-selection method (i.e. Lasso) and successively applied the appropriate regression on the selected predictors for each response (see method section for Statistical analysis). Having selected the relevant predictors for each response, we ran the logistic ordinal regression model to understand drivers for disease severity which indicated that the Shannon diversity of various SCs of *Xp* present in the field (t value = 4.15, *p* < 0.05) strongly influenced BLS disease severity as suggested in the earlier result (S6A1 Fig). Among the climatic factors, the standard deviation of Wet Bulb Temperature at 2 meters (t-value = 2.06, *p* < 0.05) and the standard deviation of the average of the wind direction at 10 meters above the surface of the earth (t-value = 2.73, *p* < 0.05) significantly increase BLS disease severity (Tables 1A and S3A), indicating that large variation in wet bulb temperature (adiabetic saturation temperature) and wind direction values led to higher disease severity. Next, a beta regression model predicting the drivers of *Xp* abundance identified one key predictor: the standard deviation of wind direction at 10 meters above the earth’s surface. This variable has a positive association with the absolute (z-value = 2.08, *p* < 0.05)) (Tables 1B and S3B) and relative (z-value = 2.69, *p* < 0.05) abundance of *Xp* (S3C Table). Additionally, kurtosis and skewness of certain climatic parameters were observed as significant, suggesting extreme and frequent shifts of climatic variables influence disease severity and pathogen abundance (S3 Table). The complete analysis providing information about other significant climatic variables to explain disease severity and pathogen abundance is available in the supplementary text (Supplementary Text).

**Table 1.** Various climatic factors influence disease dynamics and *Xp* abundance. (A) Significant climatic variables from the ordinal regression model predict the degree of correlation between various environmental factors and BLS disease severity. **(B)** Significant climatic variables from the beta regression model, showing the degree of correlation between various environmental variables and the absolute abundance of *Xp.* The Value represents the estimated increase in the log odds of the outcome for a one-unit increase in the predictor variable. The Std. Error measures the precision of this estimate. The t/z-value indicates that the effect of the predictor variable is statistically significant if the values exceed 2 or fall below -2.

| Table 1A |  |  |  |  |
| --- | --- | --- | --- | --- |
| Factors | Value | Std. Error | t-value |  |
| Shannon diversity | 2.37409 | 0.57175 | 4.1523 |  |
| Standard deviation of Wet Bulb Temperature at 2 Meters | 0.96797 | 0.46843 | 2.0664 |  |
| Standard deviation of Wind Direction at 10 Meters | 0.09547 | 0.03492 | 2.7338 |  |
| Skewness of Clear Sky Surface PAR Total | -0.94701 | 0.36778 | -2.5749 |  |
| Kurtosis of Temperature at 2 Meters Range (C) | -1.25559 | 0.59055 | -2.1262 |  |
| Table 1B |  |  |  |  |
| Factors | Estimate | Std. Error | z-value | Pr(>\|z\|) |
| Standard deviation of Wind Direction at 10 Meters | 0.03934 | 0.01885 | 2.087 | 0.0368 |
| Skewness of Surface Pressure | -1.70069 | 0.52146 | -3.261 | 0.0011 |
| Kurtosis of Relative humidity at 2 Meters (%) | -0.66357 | 0.26293 | -2.524 | 0.0116 |

Finally, we used Dirichlet compositional regression to evaluate the influence of climatic factors in explaining the differential dynamics of pathogen lineages or SCs across different fields. This type of regression ensures that dependence between cluster abundance is accounted for (i.e. the proportion of clusters must sum to one). We found that the variables that appeared to be significant across pathogen lineages were the standard deviation of specific humidity at 2 meters and mid- season (compared to end-season). In particular, these appeared to diminish abundance across all clusters except for SC6 (S4 Table). These findings suggest that frequent shifts in the climatic conditions can influence individual lineages and their abundance. Specific variables that impact individual lineages are discussed in detail in the supplementary text (Supplementary Text).

### Pathogen heterogeneity and variable fitness levels influence disease dynamics

We observed differential strain dynamics with spatial and temporal variations in dominant lineages (Fig 2C). The variation in strain dynamics can result from variability in fitness levels of different pathogen lineages and/or variable effects of environment on host-pathogen genotype interactions. This variation in fitness levels can stem from differential virulence and transmission capabilities or direct or indirect (host-mediated) competition among co-occurring pathogen lineages. We asked how pathogen heterogeneity and variable dynamics could contribute to the explanation of fluctuating disease pressures during outbreaks. We used complementary data from the greenhouse experiment and field data provided here to evaluate the hypothesis that different disease pressures are caused by different fitness levels of co-existing pathogen lineages. The greenhouse experiment involved infecting tomato plants with individual strains representative of different lineages or in mixed infections. Overall, mixed infections led to higher disease severity than individual infections and individual lineages differed in their disease severity levels (S7 Fig). For example, disease severity level of mixed infection of SC3 and SC4 was significantly higher than that of individual infections (S7B Fig).

Next, we used field metagenome data with the relative abundance of pathogen lineages and disease severity data to assess pathogen fitness under the field condition. We used StrainRanking, a regression-based method for ranking of pathogen strains with respect to their contribution to epidemics. This allowed us to combine pathogen genetic data with epidemiological monitoring of disease progression to investigate differences among pathogen lineages in their performance in the field. The fitness of an SC is the contribution of SC to the variation in the observed disease severity between the two sampling points, for example, mid and end season in this study, and is measured by a parameter estimated with the StrainRanking approach. A positive value for this parameter indicates that corresponding SC leads to an increase in the disease severity, and a negative value corresponds to a decrease of the disease severity. Lineages SC4 and SC6 tend to have a higher fitness than SC3 and other low frequency SCs with respect to their contribution to the field epidemics, as indicated by their positive contribution towards disease severity from mid to end season in the theoretical epidemic curves (Fig 3A). Equal weighted means of the genetic frequencies observed during the mid- and end-seasons (p=0.5 for each) were taken into consideration for this analysis. Even considering the weightage of genetic frequencies high (p=0.9) at the mid-season (S8A Fig) or at the end of the season (p=0.10) (S9A Fig), SC6 and SC4 contributed significantly in terms of fitness towards the growth of disease severity. Because SC6 was not predominant lineage across our samples, so we did not consider SC6 for further analysis. When strain interactions were evaluated for their fitness contributions, lineages SC4 (>20% abundance) and SC3 (≤20% abundance) combination showed positive fitness, whereas interaction corresponding to SC4 (≤20% abundance) whatever SC3 frequency led to negative fitness contribution. When SC4 (>20% abundance) and SC3 (>20%) coexisted, we observed near zero fitness contribution (Figs 3B, S8B and S9B). Next, we evaluated fitness levels of individual SCs when strain-environment interactions were considered. It’s interesting to note that we found that various contextual factors significantly impacted SC4 fitness levels but less in SC3. When average surface pressure and standard deviation of wind speed were less than mean values, but average temperature and skewness of wet bulb temperature were more than mean values, SC4 demonstrated a larger positive fitness contribution (Fig 3C). We found that specific environmental conditions that are observed in the middle or end of the season have different effects on the fitness contributions of SC3 and SC4, particularly when taking into account genetic frequencies that are reported at the end of the season (p=0.10) (S9C Fig). While lower values of the parameters promote better fitness of SC3, higher values of the clear sky PAR standard deviation, kurtosis of surface pressure, and standard deviation of temperature at 2 meters support higher fitness of SC4. Conversely, lower levels of average clear sky PAR favor SC4 fitness (S9C Fig), whereas higher values enhance SC3 fitness. However, when genetic frequencies which are reported in the mid of the season were taken into account, SC4 was more fit in the across different environmental parameters than SC3 (S8C Fig).

**Fig 3.**
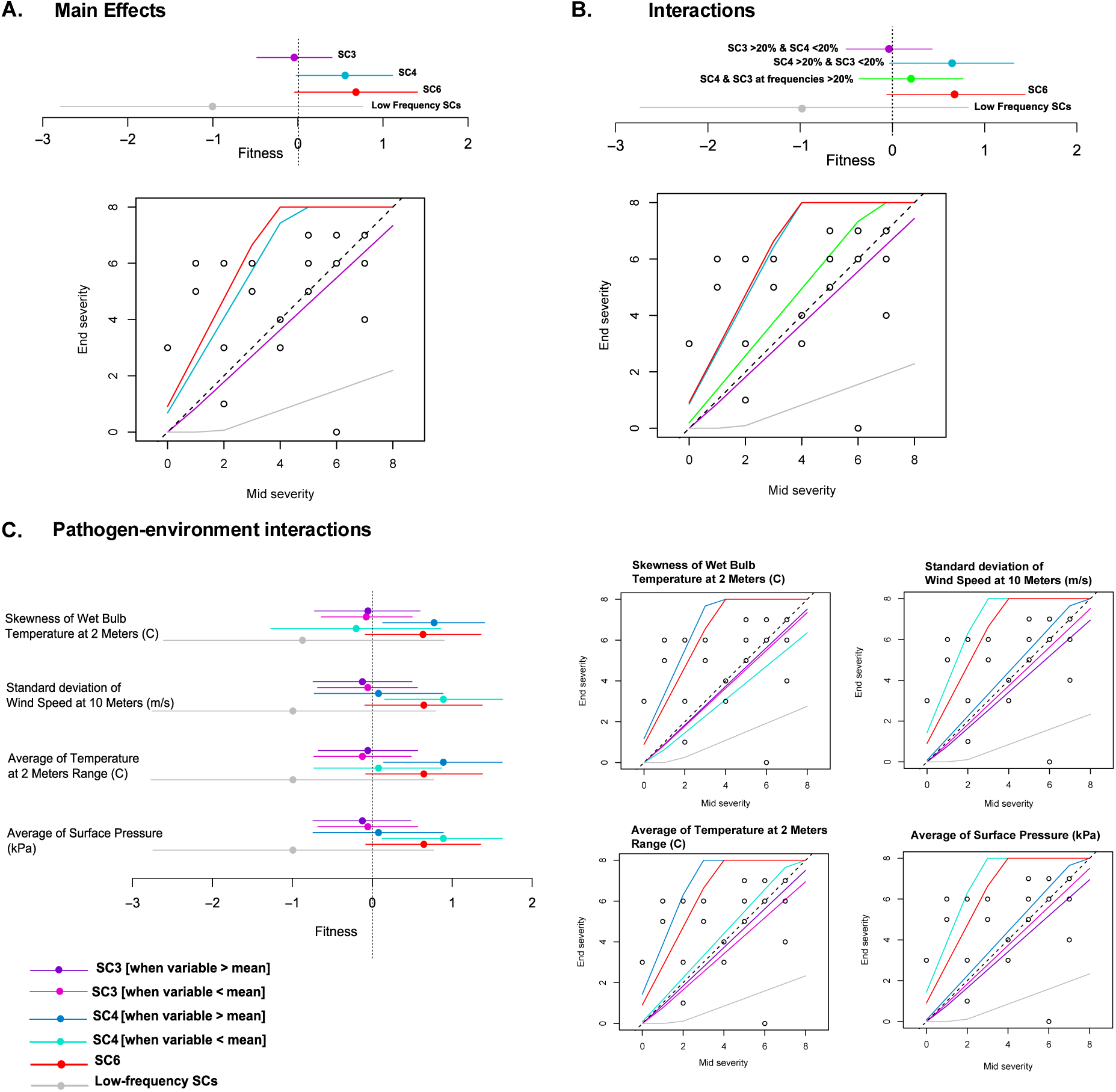
Different lineages have variable fitness levels and resilience towards different climatic factors and competition with co-occurring lineages. Using StrainRanking **(A)** Comparisons of different SCs with fitness estimates and their confidence intervals under main effects were made. These estimates focused on SC3, SC4, and SC6 (which were more abundant in samples), and a group of SCs (SC1, SC2, SC5, SC7, and SC8) that were categorized as low frequency (due to their lower relative abundance, as estimated in Fig. 2C). The bottom plot illustrates the growth curves of these SCs, estimating their contributions to the progression of disease severity from mid and end of the summer season. **(B)** Pathogen-pathogen interactions, specifically between the two dominant SCs (SC3 and SC4), were analyzed based on their combined frequencies being either below or above 20%. Fitness estimates with confidence intervals were compared when the summed frequencies of SC3 and SC4 were taken into account. Different interaction scenarios were considered, such as SC3 > 20% & SC4 < 20%, SC4 > 20% & SC3 < 20%, and both SC3 & SC4 at equal frequencies, along with SC6 and the low-frequency SCs. The growth curves in the bottom plot highlight how these interactions contributed to the progression of disease during the season. **(C)** Pathogen-environment interactions were examined by applying the frequencies of SC3 or SC4 to different environmental variables, depending on whether each variable was above or below its mean at the sampling point. The analysis assumed an equal-weighted mean of genetic frequencies to calculate fitness levels: *p* x mid-frequency + (1 - *p*) x end-frequency, where (*p* = 0.5) in this figure. All the colour keys are specific in each section of the figure and a similar colour scheme is followed for their corresponding growth curve plots.

### Tracing the allelic frequencies across seasons suggested presence of conditional SNPs that oscillate across temporal scales

While pathogen diversity analysis based on globally known pathogen lineages revealed strain dynamics driven by climatic factors and variable strain-fitness levels, we next investigated the extent of genetic polymorphism in the pathogen population (without having a bias of known pathogen lineages) by tracing allelic frequencies within and across the fields. For this analysis, a pangenome reference was constructed for *X. perforans* and allelic frequencies for different sites within the genome were obtained by mapping metagenomic reads to the pangenome. These allelic frequencies were monitored within and across seasons and fields to study the maintenance of genetic polymorphism and allele frequency dynamics. The influence of season and year on pathogen diversity led us to identify variants selected across pathogen populations over time and seasons. We looked for alleles whose frequencies changed within and across seasons and parallel across fields. For this analysis, we grouped all samples from each respective sampling time (i.e., summer or fall of a specific year) into a single metapopulation (Fig 4A). We then tracked the major and minor allele frequencies across three sampling points over three years. We observed a strong influence of seasonality on the allelic frequencies. Allelic frequencies at most sites differed across the mid/end of the summer and fall seasons. For example, several of the alleles (with *f* >=0.8) found in the middle or end of the summer growing season showed a significant change in frequency (with *f* <= 0.2) during the fall seasons for both the years of 2020 and 2022 (Fig 4B). Similarly, the alleles with their frequencies less than 0.8 during mid- 2020 remained in the population at intermediate frequencies during end seasons or mid-seasons in the subsequent years but changed to either 1 or 0 during the fall seasons of the 2020 and 2022 (Figs 4C and S10). These polymorphisms that undergo dramatic rapid shifts in allelic frequency and oscillate asynchronously between seasons repeatedly over subsequent years can be defined as conditional or seasonal SNPs. These seasonal SNPs would most likely be adaptively responding to selection pressures that vary across seasons.

**Fig 4.**
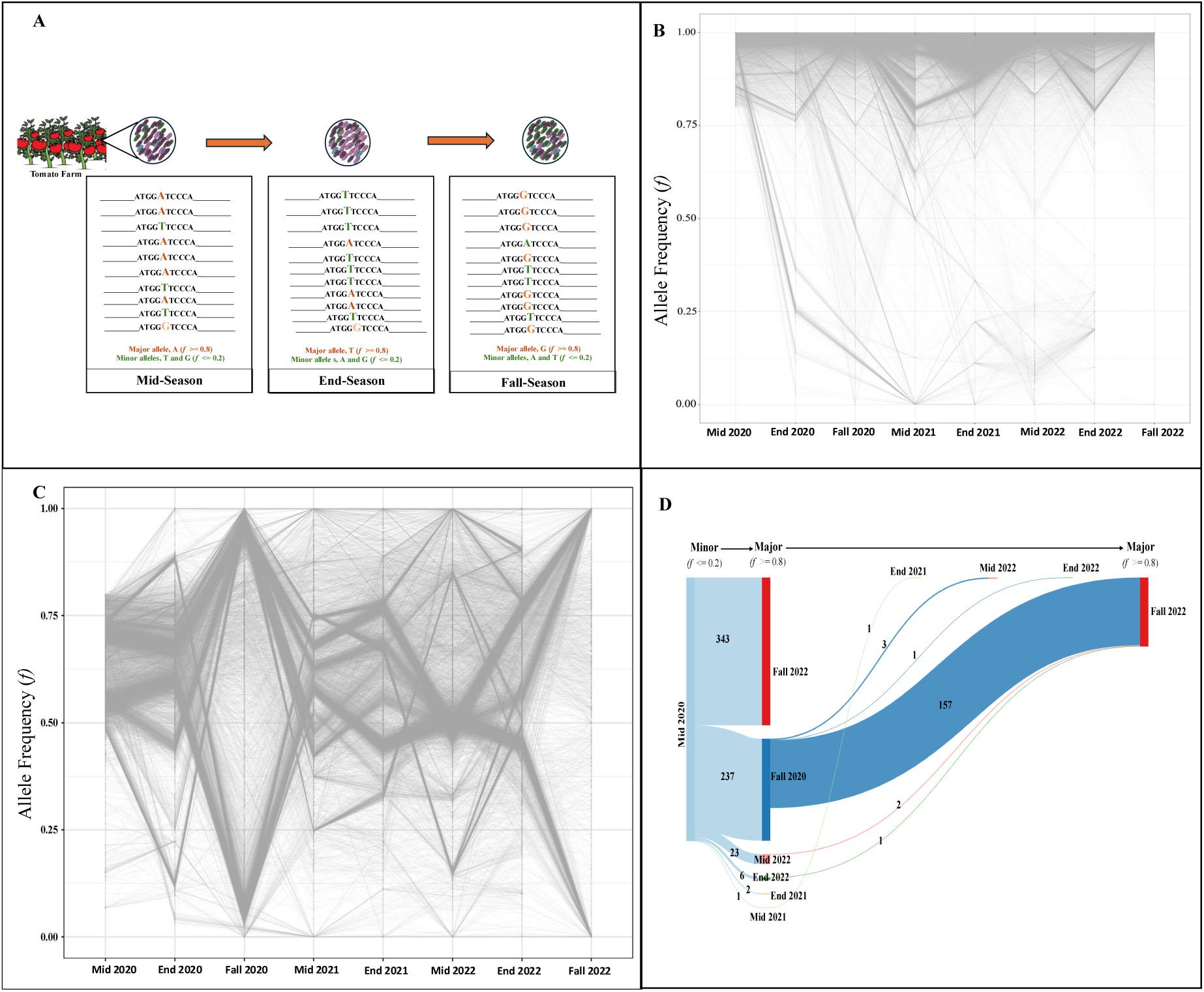
Seasonal oscillations of allelic frequencies with the signature of parallel evolution. **(A)** A graphical representation of seasonal variation, where during mid-season (of summer), allele A was the major allele (allele frequency > 0.8) in the population. By the end of the season (of summer), the A was replaced with T, and allele A became minor in population. However, by the fall season, allele G became a major allele, and the rest were present in low abundance in the pathogen population. The line plot shows alleles with their average frequencies of **(B)** *f* >= 0.8 & **(C)** *f* < 0.8 in the pathogen population during mid- season (of summer) of year 2020. The allele frequencies were tracked for different sites of *Xp* during the following seasons over three years. **(D)** The Sankey plot presenting the counts of those alleles which were present in parallel across at least 50% of the farms sampled during the mid-season (of summer) of 2020 with frequency <= 0.2 (with read depth of at least 10) and became major allele (*f* >= 0.8) in the population in the next season and counted those same alleles which stayed as major allele during the next seasons of following years.

Given the influence of seasonality on oscillating patterns of polymorphisms, we focused on identifying alleles that were seen in parallel across at least 50% of farms with *f* <= 0.2 (called a minor) during the first season and became *f* >= 0.8 (considered a major) in the following season (Figs 4A and 4C). We found that 237 of the alleles that were minor/low-frequency alleles in mid- 2020 turned major (i.e., reached high frequency) during fall 2020, with 157 of those alleles reappearing as major during fall 2022 but were not observed as high-frequency alleles in any other season of the following years. Similarly, 1234 minor alleles that existed at the end of Summer 2020 reached high frequency (i.e., major alleles) in the fall of 2020. Of these, 138 alleles were observed at high frequency in the subsequent fall season of 2022 (S11A Fig). Furthermore, we observed 3598 allelic sites with allele frequency less than 0.2 during the fall of 2020 became major during mid-season (summer) of the year 2021 and most of the sites (1025) appeared back as major alleles in the mid-season of the year 2022 (S11B Fig).

### Despite variability in disease dynamics, signatures of selection and parallel evolution were evident in the pathogen population

Given the varying disease pressure observed across different farms, we sought to determine if any genes in the pathogen genome were consistently under selection across these locations. Using Metapop package, we estimated the pN/pS (non-synonymous to synonymous ratios) of different genes of *Xp* and pooled out the genes with pN/pS > 1 values. We filtered out the genes if they were present in at least 50 % of farms in a particular sampling point to provide evidence for parallel evolution. We found eight genes under selection were parallel across farms (at least 50 % of farms of specific sampling time) and seasons. These genes comprised type VI secretion system (T6SS) protein ImpG, ClpV1 family T6SS ATPase, S9 family peptidase, aminotransferase, tonB-dependent receptors, S9 family proteins (S12 Fig and S5 Table).

However, none of these eight genes were clustered and are present at distinct locations based on the reference genome of *Xp* LH3. Interestingly, two proteins associated with the T6SS were under positive selection, suggesting the microbiome’s potential role in selection pressure and shaping the pathogen genome. We also identified a gene, class 1 fructose-bisphosphatase, under positive selection only in the populations from both fall seasons of 2020 and 2022 (S5 Table). Overall, these results suggest evidence of parallel evolution across farms and seasons in the *Xp* pathogen population.

## Discussion

The significance of strain diversity, strain fitness and compositional dynamics towards disease epidemics has been highlighted in wild plant pathogens but the importance of genetic polymorphism in explaining variability in disease outbreaks is less understood in the agricultural settings, partly due to an assumption of genetically monomorphic populations under monoculture system and the emphasis on the use of methods that lack resolution of pathogen population structure at microgeographic level. In this work, we developed an approach to study the extent of genetic polymorphism in the pathogen population within individual fields over the course of a growing season and over three years, linked the pathogen dynamics to climatic variables to explain variability of disease dynamics in the fields in neighboring states (Fig. 5). We found that as many as five pathogen lineages can co-exist within a single agricultural field. Prior studies have documented that under natural systems, no single lineage takes over and multiple lineages co-exist at low abundance but in contrast, single dominant lineage is assumed to dominate in the agricultural fields (25). We found that at least three distinct lineages dominate across different fields and at times, two distantly related lineages can co-exist within an individual field at high abundance throughout the growing season (Fig 2). The differential strain dynamics was observed across neighboring fields and was linked to differential climate sensitivity of the lineages and differential fitness contributions towards epidemics (Fig 3). When we conducted analysis of pathogen dynamics without having bias of known lineages, but by tracing the allelic variants over three years, the analysis revealed a considerable influence of seasonality on pathogen dynamics, with distinct seasonal alleles strongly favored throughout specific seasons in the consecutive years (Fig 4). This genetic heterogeneity with differential and environmentally dependent fitness contributions might be an adaptation strategy that enables pathogen to dynamically respond to environmental fluctuations within seasonal timescales. These complex interdependent pathogen genotype-genotype-environment interactions may complicate the selective pressures during a host-pathogen arms race in case of endemic diseases in agricultural systems. Understanding the roles of conditional seasonal alleles that we observed in this work and the role of loci under selection in dictating the pathogen fitness represent an important avenue for future work. Identifying the drivers of the disease dynamics and pathogen diversification will be critical in reducing the frequency of pathogen outbreaks (Fig 5).

**Fig 5.**
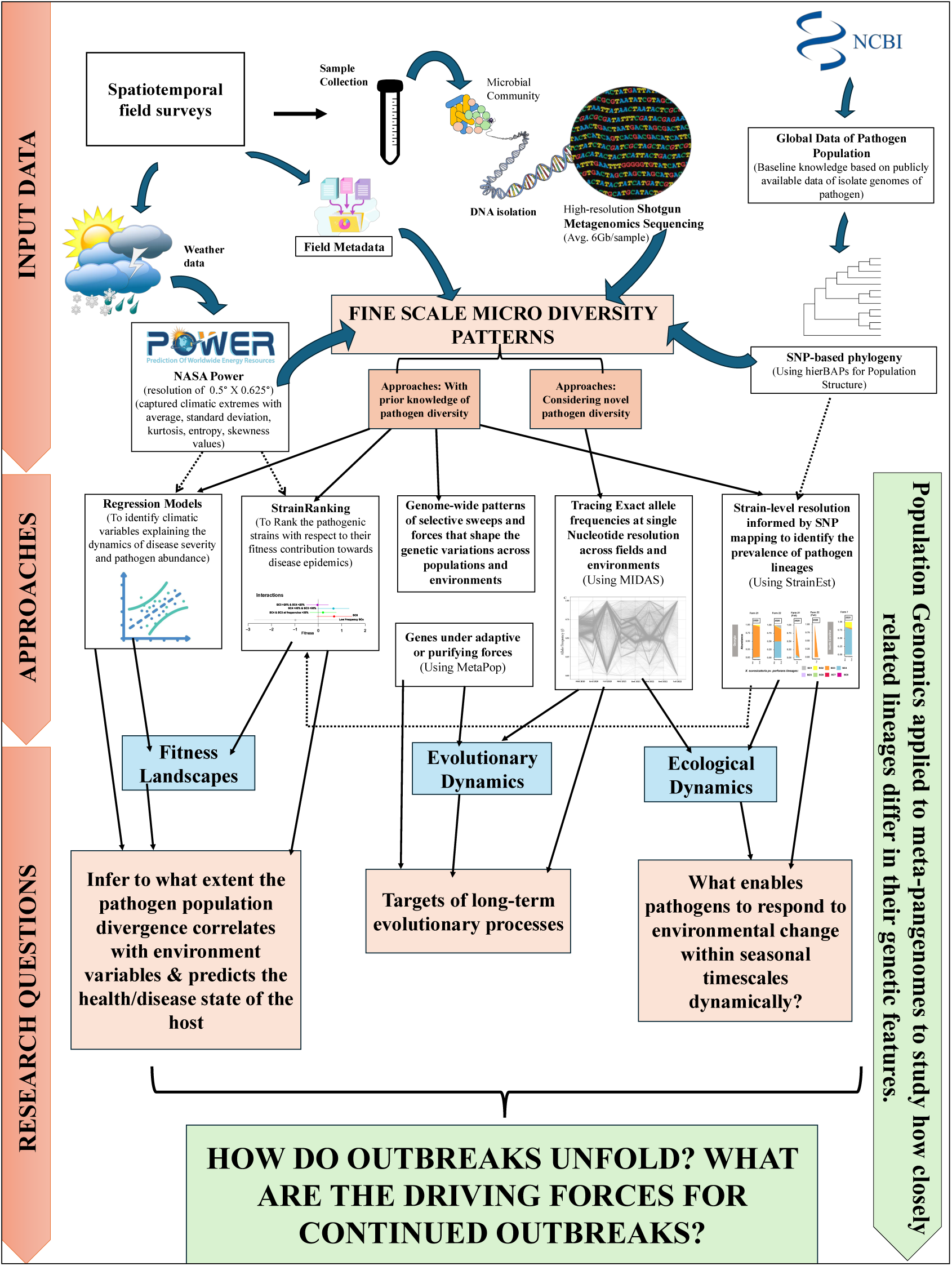
Integrated workflow to study pathogen outbreaks incorporating large scale field survey data, high-resolution metagenomics data and subsequent analyses to understand eco-evolutionary pathogen dynamics.

Regression models applied to link environmental data, disease outcomes and pathogen diversity in this study revealed that extreme climatic events were significant drivers of disease dynamics and pathogen diversity across different farms. More variations and extreme events such as changes in wind direction, photosynthetically active radiation, temperature, humidity, and surface pressure (Tables 1 and S3) can directly influence pathogen dispersal (52–54) and plant host physiology (55–57), or indirectly alter host-pathogen, strain-strain, and pathogen-microbiome interactions (39,58,59). While empirical evidence for how recent climatic shifts have contributed to disease outbreaks is limited, studies on the impact of extreme weather events on the virulence of fungal pathogens like *Hemileia vastatrix* (coffee rust), *Puccinia striiformis* (wheat stripe rust), and *Phytophthora cinnamomi* (Mediterranean oak decline) have highlighted the need for these studies (60–62). Our approach of capturing the climatic extremes using measures such as standard deviation, kurtosis, skewness of climatic variables observed for each farm across seasonal timescales allowed us to assess the extent to which pathogen genetic divergence correlates with climatic shifts. The findings from these models highlighted the importance of capturing these climatic extremes in the epidemiological modeling, rather than average climatic variables. For example, parameters that we identified as significant (negative impact on disease severity and pathogen abundance) were more extreme values of temperature and humidity. Our observation of more variation in wind direction being significantly associated with higher disease severity and higher pathogen abundance align with previous observation of importance of dispersal in the epidemic growth for bacterial spot disease. The aerosolization and long-distance dispersal of <=2m was shown for this pathogen under humid transplant house conditions (63). Later a single effector, XopJ2, was shown to provide increased fitness by increased dispersal in the field (64).

Our findings of co-existence also challenge the traditional view that a single lineage of an abundant pathogen dominates the host population in agricultural systems. Two lineages, SC3 and SC4 co-existed, both at large densities, but neither showed a strong dominance across different fields in all three examined growing seasons (Figs 2C, S4). First, this observation is intriguing, because it certainly challenges the competitive exclusion principle, which states that two species occupying the same ecological niche cannot coexist indefinitely (65). Second, it reveals an apparent paradox because, based on our analyses, SC4 seems to be fitter than SC3. SC6 also seems to be as fit as SC4 but is present at a lower level with distribution limited to only certain fields in Alabama than SC3 and SC4, thus, its contribution in the lineage dynamics could not be systematically assessed. Here, we sought explanations for this apparently unexpected co-existence of SC3 and SC4 and their ecological dynamics. Prior studies on *E. coli* have demonstrated the prevalence and persistence of season-specific strains due to fluctuating temperatures throughout the year (66–68). However, we have yet to observe any clear patterns of early or late-season specialization of individual SCs, indicating that time partitioning is unlikely to explain their co- existence. Since we did not sample outside the host, current data cannot explain whether these lineages have high fitness outside the host or neighboring plant species (non-host). In addition, lack of data on host genotypes or field landscapes do not allow to evaluate whether host heterogeneity and landscape heterogeneity in the field supports their co-existence (44). Another limitation of our approach which used pooled leaf samples representative of a field for metagenome sequencing, was that we couldn’t exclude spatial partitioning or microhabitat difference (for example, leaves of different ages, or patchy distributions within individual fields) that can support co-existence of two or more pathogen lineages. Resource specialization cannot be ruled out either because the dominant lineages coexisting in the fields, SC3 and SC4, differ in type III effector content and may differ in within-host multiplication rate and transmission rate, or they may prefer different niches, such as leaf surface vs apoplast. Our finding that sudden seasonal events or climatic fluctuations had a significant impact on defining individual lineages’ fitness contributions may potentially explain their coexistence. Two independent approaches assessing compositional dynamics (S4 Table) and fitness dynamics highlighted the influence of environmental parameters on individual lineages (Figs 3, S8 and S9), specifically SC3 and SC4, specifically climatic fluctuations as indicated by skewness, kurtosis and standard deviation of different parameters, being significant in dictating abundance of SC3 or SC4. We observed that field-level fitness contribution by SC4 seemed to be impacted by other biotic (SC3) and abiotic factors, whereas that by SC3 did not. The varying performance of SC4 depending on different environmental conditions (from very fit to intermediately fit: fitness = 0) may contribute to the similarity of its overall frequency with the SC3 frequency (more generalist with respect to the values of tested factors and the presence of other SC). Thus, maintaining co-existence of lineages that show overall environmental resilience as well as environmentally dependent fitness might be a strategy that pathogen adopts to cope up with the sudden climatic shifts. All the arguments made above that may contribute to explain the co-existence of SC3 and SC4 at relatively high and similar levels, and the estimation of their relative fitness, can be summarized by making a parallel with the effective reproduction number, Rt, classically considered in human and animal epidemiology. Rt is the average number of new infections caused by a single infected individual at time t in the partially susceptible population. Rt varies in time because of the variation of the number of susceptible individuals but may also vary because of temporal heterogeneity in the eco-evo- determinants of the transmission (e.g., because of lockdown measures during the covid-19 pandemic; (69)). Rt can obviously be considered at the lineage level. Broadly, the epidemic or the lineage dynamics tends to keep going on if Rt ≥ 1. In terms of fitness, as estimated by the StrainRanking analysis in this study, it is equivalent to a fitness value ≥ 0 (the fitness here is an indicator that integrates diverse processes and variables and is therefore an aggregated measurement of the capacity of the lineage to “multiply” in real conditions). Thus, SC4 with fitness >0 and SC3 with fitness of around 0 indicate Rt ≥ 1, suggesting that both SC3 and SC4 are fit to keep epidemic going or to expand. As Rt evolves in time, the fitness (viewed as an aggregated measurement) may also evolve in time, and an interesting research area would be to finely characterize the eventual variation in time of the SC’s fitness, evaluation of lineage-specific genes in contribution towards fitness, and to determine how the possible processes evoked above contribute to this variation. This is because possibility of transitory state with co-existence as opposed to equilibrium state cannot be ruled out, especially, given the limited geographical region studied here (70) and that genetic divergence has been observed in this pathosystem (Potnis et al. 2021). Thus, continuing such high-resolution lineage fitness tracking across tomato producing regions and disease outcomes will be important outputs to predict future outbreaks, and to design disease management strategies that are tailored to concurrent pathogen population structure (i.e. breeding for disease resistance and climate resilience).

Although seasonality was not evident at the individual lineage level, analyzing the data via variant tracing indicated the presence of season-specific allelic pathogen variants (Fig 4), particularly unique to the summer vs fall growing season over the three consecutive years of sampling (Fig 5). These seasonal variants might be indicative of a trade-off between in-season within-host multiplication, in-season transmission, and inter-season survival, where variants that contribute as primary inoculum (the form under which pathogen survives host absence) in fall vs summer might differ due to the variable inter-season survival periods experienced (71). One approach to test this notion is to see if there is a trade-off between survival outside the host (with varied overwintering durations) and infection efficiency (72). Periodic host absence has been claimed to cause evolutionary divergence (also known as evolutionary branching), however, contradictory evidence exists (73–75). The absence of year-round production in Alabama, South Carolina, North Carolina, and Georgia may drive evolutionary divergence. The seasonality could also mean differing day lengths altering host physiological responses to the pathogen, thus impacting the fitness contributions of different variants towards epidemics (76) or differing light intensities impacting pathogen behavior and virulence, as seen with differing motilities and differing epiphytic fitness in the case of *Pseudomonas syringae* strains (77). The findings of seasonal oscillations of genome-wide allelic frequencies may suggest evidence of fluctuation selection contributing towards the genetic variation observed in this population.

We also looked at signatures of parallel evolution by identifying loci under positive selection across > 50% of the fields (S5 Table). Notably, we identified certain genes under positive selection over three years, including the TonB-dependent receptor gene located adjacent to the PepSy domain-containing protein. The widespread use of heavy fertilizers and pesticides in agricultural systems (such as mancozeb applications to manage bacterial spot contain Mn and Zn) may have been responsible for the selection of this TonB-dependent receptor likely involved in heavy metal transport (78,79). Furthermore, multiple genes belonging to two separate Type VI secretion system clusters were found to be under selection, suggesting that the microbiome may play a significant role in driving the selection within the pathogen genome (80–82).

Consistent with the assumptions of seedborne diseases and their global spread (30,83–85) in the agricultural fields, our survey indicated that almost all known *X. perforans* lineages identified in the global population are circulating in the southeastern US fields. Although multiple lineages co-existed in the individual fields, no such apparent link between genetic relatedness of co-occurring lineages and disease severity was observed, as was previously shown to be important through the mechanism of competition or increased virulence (86). This could be because co-occurrence of SC3 and SC4 was predominant in majority of the cases, with other low frequency lineages not particularly contributing towards fitness and we lacked consistent sampling of the fields throughout the growing season. However, higher diversity (Shannon as well as average nucleotide diversity) was correlated with higher disease severity (S6 Fig). Such observation could be explained by ability of diverse strains to manipulate the host defenses in a variety of ways (87,88), or through resource-mediated competition or through sharing of public goods, such as quorum sensing signals (89). While our survey collected information on host cultivars, the lack of information on host genetics or levels of tolerance/susceptibility did not allow evaluation of the role of host genotypes in driving disease dynamics. However, age-related host resistance (90–92) or the prevalence of other diseases later in the season might contribute towards seasonal disease pressure and thus, seasonal dynamics that we observed. The regression analysis also included scale of the farm as one of variables. While disease severity was not significantly different across small vs commercial farms, more pathogen diversity was observed in the commercial farms compared to small farms (S3 Fig). This intriguing observation brings about apparent contradiction, i.e. small-scale farms often have transplants, seeds from a variety of sources, thus, possibility of introduction of multiple pathogen genotypes. However, crop/genotype diversity in the small-scale farms with intercropping may also lead to reduction in pathogen abundance and diversity (93). These observations demand further investigation of whether such intercropping might be a resilient strategy for disease control.

Taken together, we find that plant-pathogen interactions in agricultural environments are much more complex than previously thought. Our study revealed that maintaining pathogen heterogeneity and variable fitness with seasonal allele fluctuations in response to extreme weather events are the strategies employed by the pathogen. This work also presents empirical evidence on the influence of recent climatic changes on altering disease outcomes. We suggest that breeding programs need to consider multiple pathogen lineages that show positive fitness contributions based on large scale field surveys. Tracing these evolutionary changes can predict which lineages will be a better fit in the long run, especially in the face of sudden climatic shifts, aiding in developing more effective breeding strategies. Future work based on the results of this study should focus on identifying the roles of specific fitness factors in pathogen biology, as this could provide valuable insights into pathogen adaptation strategies and the selection pressures they face. Finally, we present an integrated workflow utilizing metapangenomics to infer fine scale microdiversity patterns in the pathogen population, coupled with epidemiological and statistical modeling to link climatic variables to the eco-evolutionary processes impacting disease risk and to study disease epidemics at large scales (Fig 5). This approach can allow disentangling targets of long-term eco-evolutionary processes and for studying fundamental principles involved in response of microbes in dynamically responding to environmental shifts. Such approach can be applied to different case studies in plant, animal, and human health.

## Materials and Methods

### Reconstruction of the *X. perforans* core genome using publicly available *Xp* genomes from NCBI

Sequencing reads for 467 *Xp* strains downloaded from NCBI (as of 08/15/2023) were used to study the global diversity of *Xp* isolate genomes. To ensure high-quality and minimally biased genomes, the genomes were de-replicated using dRep (v3.2.2) (94) using a 95% minimum genome completeness and 5% maximum contamination. We manually discarded the genomes with more than 400 contigs to ensure better alignment with the reference genome and avoid bias while creating the SNV profile. Four hundred sixty-seven quality-controlled genomes collected from tomato and pepper (with one eggplant isolate) around the globe were individually aligned against the completed genome of *Xp* strain LH3. Single-nucleotide polymorphisms (SNPs) common to all genomes were extracted to generate a concatenated set of high-quality core-genome SNPs using Parsnp (95). The phylogenetic tree was visualized and annotated using iTol (v.6.7.5) (96)

### Sample collection and disease severity estimation

To study the diversity of BLS *Xanthomonas* in the southeastern United States, we collected 66 leaf samples from 22 fields, designated as either commercial-scale or small-scale (Table S1). These samples were collected from symptomatic and asymptomatic tomato plants grown in Alabama, Georgia, South Carolina, and North Carolina over 2020, 2021, and 2022 (Fig. 1A). Samples were taken during mid-season, end-season of the summer growing season, and when available, in the fall growing season (S1A Table) to capture the temporal dynamics of the pathogen population. While sampling, two to three leaflets were randomly taken from individual plants in the field using variable sampling methods based on the scale of the field (Fig. 2B). Multiple plants were sampled across the field to obtain a representative 50-100 grams leaf tissue sample. Assessments of the disease were made by estimating the percent of disease symptoms and defoliation caused by bacterial spots (S1B Table) using the Horsfall-Barratt scale, which ranges from 1 to 12, with scale of 1 being no disease and 12 being 100% defoliation (51).

### DNA extraction and sequencing

Metagenomic DNA extraction, quantification, and sequencing of the samples were done as described earlier (23). Approximately 40 grams of the leaf sample was taken in a Ziploc bag (Ziploc®) with 50 ml 0.1% wash buffer (0.05M PBS, 8.5 g NaCl, and 0.2 ml tween 20 per liter of water), sonicated for 20 minutes, followed by transfer of the buffer to a 50 ml falcon tube with the help of a sterile plastic serological pipette. The tubes were centrifuged at 4000 rpm, 4^°^C for 20 minutes (Eppendorf^®^ 5418 R) to collect all the cells. The cells were then washed twice with sterile deionized water, and the metagenomic DNA extraction was done using Wizard® Genomic DNA Purification Kit (Promega) as per the manufacturer’s instructions with the addition of a phenol-chloroform step to remove protein contamination. The extracted DNA was quantified using a Qubit^®^ fluorometer and dsDNA high-sensitivity assay kit (Life Technologies, Carlsbad, CA, USA) and kept at -80^0^C until further processing. Sequencing was performed on the Illumina HiSeq 4000 platform. The raw reads were processed and trimmed for the adapter using BBDuk (Bushnell, 2014). DynamicTrim command of SolexaQa was used to trim the reads based on quality values with Qphred less than 20, followed by filtering the reads shorter than 50bp (97). Host reads were separated and removed using tomato cv. Heinz 1706 as a reference (GCA_022405115.1) from the metagenomic samples using KneadData tools (98) as a reference.

### Taxonomic profiling and diversity estimation

The relative abundance of *Xanthomonas* sp. in quality-controlled and host-decontaminated reads was performed using Kraken2 (v2.1.2) (99) against a standard Kraken2 standard database. The database contained RefSeq libraries of archaeal, bacterial, human, and viral sequences as of March 1, 2022 (100). The resulting Kraken2 report files were used as inputs for Bayesian re-estimation of abundance using Kraken (Bracken) (v2.6.2) (101). We also calculated the absolute abundance estimates for *Xp* by combining their relative abundance with the total DNA content in each sample. Microbiota density, expressed as total DNA (ng) per mg of fresh sample, was computed for each sample. This information was utilized to determine the absolute abundance of *Xp* by multiplying the amount of DNA per mg of the sample with its relative abundance.

To explore the intraspecific diversity and dynamics of *Xp* in the metagenomic samples, we employed the StrainEst pipeline (102). StrainEst is a reference-based method that utilizes single-nucleotide variant (SNV) profiles from carefully selected reference genomes. This approach enables us to estimate strains’ presence and relative abundance within a given sample. Initially, we conducted SNV profiling of representative strains from *Xp*, *Xeu*, and other closely related pathovars belonging to *Xeu* species complex to determine their presence. A second round of profiling was done to identify the diversity of *Xp* strains. A representative genome from each *Xp* SC (Fig 2) was selected and aligned against the completed reference genome of *Xp* strain 91-118 using the MUMmer algorithm (103). This genome alignment was used to construct a SNV matrix, where each row corresponded to a variable position in the reference genome, and columns contained allelic variants present in the reference strains. This matrix was used as a reference in modeling strain level abundance.

To assess the diversity of *Xp* SCs within the samples, we computed the Shannon diversity index (Shannon-Wiener diversity index) using the “diversity()” function from the R package *vegan* (v2.6-4) (104). The index considers the count of SCs present (richness) and their proportional distribution (evenness). A higher index value indicates greater species diversity in the habitat. When the Shannon diversity index of a sample is equal to 0, it implies that only a single SC is detected in that sample.

### Weather data

Daily climate data spanning three years for all 22 fields were compiled from the NASA Power database using the nasapower (v4.0.12) package (105) in R. Data were collected during mid-season, end-season of summer, and fall samples as per the sampling time in Table S1. The gridded weather data used in this analysis are based on satellite observations and models and are offered globally at a horizontal resolution of 0.5° X 0.625° latitude–longitude grid cell. Weather variables, such as temperature, precipitation, solar radiation, relative humidity, wind, and others (S2 Table), were consolidated over time and space. The intervening period between the planting and sampling times was accounted for in the analysis for each weather parameter by calculating averages. For instance, the mid-season average was determined by averaging daily weather data from the planting date to the mid-season sampling time, which usually occurred 60 days after transplanting. The end-season average was calculated from the planting date to the end-of-season sampling time, typically 30 days after the mid-season sampling. To account for interseason climatic events, variability among the weather parameters was measured using descriptive statistics (standard deviation, skewness, entropy, and kurtosis) for all the climatic variables to see if the extreme weather events during the growing season impact disease severity, pathogen diversity, and abundance. Standard deviation measures the deviation from the mean value. A higher standard deviation indicates greater weather parameter variability, while a lower standard deviation suggests a more consistent pattern. Skewness measures the degree and direction of asymmetry. A positive skewness implies more extreme high weather parameters, while a negative skewness suggests more extreme low, with zero suggesting a symmetric distribution. Kurtosis is a measure of tail extremity reflecting the presence of outliers in a distribution (106). Higher positive kurtosis implies extreme weather values or outliers, while negative kurtosis indicates a flatter distribution with less extreme weather values. Similarly, entropy quantifies the level of unpredictability in the weather values. The higher the entropy, the more unpredictability in the weather fluctuations will be.

### Statistical analysis via regression models

We quantified the relative abundance of various *Xp* lineages across the samples and analyzed their distribution patterns in relation to different climatic factors. This analysis was designed to explain three different types of responses based on climatic and other variables: (i) relative abundance, (ii) absolute abundance, and (iii) disease severity. Before running these regressions, we chose to run a predictor-selection procedure for each of these (given the high number of predictors we considered for this work) by using Least Absolute Shrinkage and Selection Operator (Lasso) regression, which jointly aims at correctly predicting the response and selecting the most associated variables (107). As no implemented or stable versions of the Lasso directly address the types of responses we have considered in our work, we decided to use proxy responses to run the Lasso and then use the selected variables from this step to run the specific regressions and inference described above. More specifically, we use a logistic Lasso to select the variables for the (relative abundance) where we classify low and high abundance based on the 50% threshold. In contrast, we use standard Lasso regression on the severity scales (assuming they can be considered as continuous) and the Shannon entropy (a continuous scale proxy for the distribution of abundance between clusters). Predictor-selection was performed on the variables that still have non-zero coefficients after the shrinkage process using the “cv.glmnet” function to prevent the overfitting of the model. This study determined the tuning parameter for regularization using 10-fold cross-validation. The path length (min_lambda/max-lambda) was chosen to be 0.001, and 100 default values along the regularization path were tested to find the best lambda value. Once a selection was run for all responses (proxies) of interest, we ran the appropriate regressions for each response. More specifically, since disease severity is measured on a discrete ordered scale in the 0-12 range, we used an ordinal logistic regression, which accounts for the order of severity and the discrete scale of the response (108). On the other hand, as the abundance (both absolute and relative) of *Xp* is measured on a 0-1 range, we employed a Beta regression (109) using the *betareg* package (110). The beta regression model is employed for non-normally distributed data with a range between 0 and 1, addressing heteroskedasticity and asymmetry issues (109), and is specifically chosen here due to the non-normal distribution of *Xp* relative abundance in the sample, which conforms to a beta distribution. To analyze the distribution of abundance among different Xp lineages, we used a compositional regression approach, which ensures that the regression model explains and preserves the property that the total abundance among the clusters should sum to one (indeed, a separate regression for each cluster would not consider the dependence between clusters). We used the Dirichlet regression approach, using the *DirichletReg* package in R, to describe how the distribution of various *Xp* lineages changes according to changes in these potential predictors (111). As for the previous response variables, the use of the Lasso for this type of regression is computationally unstable. Therefore, we chose to use the Shannon diversity of *Xp* as the response when using the Lasso: this is a reasonable proxy since Shannon diversity is related to how the data is distributed (in this case, across the lineages). Once this was obtained, 18 variables were selected to run the Dirichlet regression.

### Greenhouse Experiment

We evaluated the effects of different SCs of *Xp* on 4-5 week old tomato cultivar FL47 plants grown under greenhouse conditions. Plants were dip-inoculated for approximately 30 seconds in bacterial suspensions containing 10⁶cfu/ml of representative strains from different SCs (SC3 (Xp2010), SC4 (AL57), SC5 (AL37), and SC6 (AL65)). These SCs were chosen based on their observed abundance in samples from our previous study (23). Treatments included a control (0.01M MgSO₄), individual inoculations with SC3, SC4, SC5, and SC6, as well as mixed co-infections of SC4 + SC3, SC5 + SC6, SC3 + SC6, SC3 + SC5, and SC3 + SC5 + SC6. All treatments were supplemented with 0.00025% Swilvet and replicated five times, with plants randomly arranged in the greenhouse. Plants were watered daily, and disease severity was assessed on days 7 and 14, with additional evaluations on days 9 and 12 for plants exhibiting delayed symptoms. Disease severity was measured using the Horsfall-Barratt scale (51). To minimize bias, two observers recorded the disease severity, and the average score was used. Raw disease severity indices were then used to calculate the area under the disease progress curve (AUDPC), which correlates multiple observations of disease progression into a single value (112). AUDPC values were plotted for each treatment across seven experimental batches using the “dplyr” package in R Studio and built linear and linear mixed-effects models using “lme4” and “lmerTest” (113,114). For individual strains a generalized linear hypothesis test was used while for multiple comparisons Tukey’s method was used. Linear mixed effect models were used to assess the development of disease severity slopes across treatments.

### Strain Ranking

StrainRanking (115) was used to estimate and compare fitness differences between SCs (main effects), between SCs considered alone or in association (pathogen-pathogen interaction), and between SCs under specific environmental conditions (pathogen-environment interaction). StrainRanking requires input data consisting of the frequencies of the SCs and the measurement of disease growth or decline for each sampling location (i.e., fields), and estimates the fitness as the relative contributions of SCs to the growth or the decline of disease severity between two sampling times (i.e., Mid and End sampling times). SC frequencies were computed as a weighted mean of Mid and End SC frequencies: *p* × Mid frequencies + (1 − *p*) × End frequencies, where *p* is equal to 0.1, 0.5 or 0.9 to consider different representativeness of Mid and End frequencies in terms of disease dynamics between Mid and End sampling times of summer season. (Because of low sample size, we did not include Fall season samples in this analysis.) Significance tests performed in StrainRanking are based on randomization, which adequately accounts for the relatively small sample sizes encountered when distributing data at the resolution of SCs or even at a higher resolution (when one considers pathogen-pathogen or pathogen-environment interactions). To account for the specificity of the severity measurement in the present study, which is a mark ranging between 0 and 8, we added to the StrainRanking method a saturation level of the growth and the decline at +8 and -8, respectively. In addition, we developed a function providing confidence intervals for estimated fitness in addition to the pairwise comparison tests embedded in the original StrainRanking R package.

For the study of main effects, StrainRanking was applied to SC3, SC4, SC6 and the group of SCs with low frequencies (SC1, 2, 5, 7, and 8), say SCLowFreq.

For the study of pathogen-pathogen interaction, only the interaction between the two dominant SCs was accounted for, namely SC3 and SC4. We consider SC3xSC4 interaction depending on whether SC3 and SC4 are lower or larger than 20%. Thus, StrainRanking was applied to the summed frequency freq(SC3)+freq(SC4) when freq(SC3)>20% and freq(SC4)>20%, and to the frequencies of SC3 when freq(SC3)≤20% or freq(SC4)≤20%, SC4 when freq(SC4)≤20% or freq(SC3)≤20%, SC6 and SCLowFreq. Not that the interaction has to be understood at the field level here since we do not have information indicating whether or not SC3 and SC4 were collected on the same plants.

For the study of pathogen-environment interaction, only the interaction with the two dominant SCs was accounted for. This step incorporated only the significant environmental parameters identified through statistical analyses using the regression model, along with their standard deviation, skewness, entropy, and kurtosis. Hence, for each factor *X* (e.g., average temperature, kurtosis of wind speed, etc.), StrainRanking was applied to the frequencies of SC3factor+, i.e. the frequency of SC3 when *X* > *X̅*(SC3) (where *X̅*(SC3) is the mean of *X* at sampling points where SC3 is present, i.e. freq(SC3)>0), SC3factor- (when *X* ≤ *X̅*(SC3)), SC4factor+ (when *X* > *X̅*(SC3)), SC4factor- (when *X* ≤ *X̅*(SC4)), SC6 and SCLowFreq. The value of *X* is the weighted mean of *X* observed at Mid and End sampling times: *p* × [*X* at Mid point] + (1 − *p*) × [*X* at End point], where *p* takes the same value as for the weighted average of genetic frequencies.

### Construction of *Xanthomonas* pan-genome

Initially, we downloaded 287 strains of *Xp (Xanthomonas euvesicatoria* pv. *perforans)* and *Xeu (Xanthomonas euvesicatoria* pv*. euvesicatoria)* from the NCBI RefSeq database (as of February 16th, 2022). Using SuperPang (v0.9.4 beta1) (116), we constructed a non-redundant pangenome as a reference for the subsequent analyses.

To avoid read stealing and donating of *Xanthomonas* with other microbiome species, we implemented an additional filter to remove genes with more than 97% similarity in our metagenomic reads. These genes were identified similarly to the method described by Kaur et al., 2024. We used the “run_midas.py species” command to identify the predominant species in our samples, utilizing the MIDAS (v1.3.2) default database as a reference, which consists of 31,007 bacterial reference genomes organized into 5,952 species groups (117). We then built a custom database of non-redundant pangenomes (*Xp* and *Xeu*) using MIDAS and subjected it to BLAST (blast+) with the concatenated gene sequences from species with a mean abundance > 0 from the previous step. Genes with 97% similarity to those in the database were removed from all samples using BBDuk (v37.36) (118). These reads were used for our subsequent analyses.

### Genes under selection

We discarded the samples with zero *Xp* abundance in the following evolutionary analyses. To calculate the genetic variation in our samples, we first mapped the samples to the non-redundant pangenome using BWA-MEM (v0.7.12) (119). Next, we removed low-quality alignments and duplicate reads using samtools (v1.11) (120) and picard (v1.79) (121), respectively. The resulting bam files were then processed with MetaPop (v1.0) (122) to calculate the ratio of non-synonymous to synonymous polymorphisms (pN/pS) for each gene. In order to find the genes under positive selection, we chose the genes with pN/pS ratios >1 and present in at least 50% of the samples in a particular season. Average nucleotide diversity values for each farm were also estimated using this package.

### Seasonal variants

To calculate the allele frequencies of SNV sites across the pangenome, we used the MIDAS “midas.py snps” script (117). We treated all samples from different states in the southeastern USA as a single population for each season. To increase the accuracy of results, we had additional filters for removing SNV sites with a read depth of less than 10. For seasonal oscillations, we averaged the allele frequencies of major alleles during the season. To identify parallel variants in the population, we first extracted SNV sites where the frequency of minor alleles was less than 0.2 (considered minor alleles) and changed to greater than 0.8 (considered major alleles) in the following season and remained major in subsequent seasons. For example, we identified sites with minor alleles (*f* <= 0.2) during mid-season 2020 that became major alleles (*f*>= 0.8) by the end of the 2020 season. We retained only those SNVs that exhibited this trend in at least 50% of the samples in a given season.

## Acknowledgments

The climatic data were obtained from the NASA Langley Research Center (LaRC) POWER Project funded through the NASA Earth Science/Applied Science Program. This work was made possible in part by a grant of high-performance computing resources and technical support from the Alabama Supercomputer Authority. We thank growers in the southeastern US for supporting this project, allowing sampling their fields, providing metadata and communicating with us their concerns in tomato production. This helped us better understand the nature of the outbreaks. We thank the Alabama Agricultural Experiment Station, and Auburn University for providing resources for conducting this work.

## Data availability

Sequence data generated from this work have been deposited in the SRA (Sequencing Read Achieve) database under the BioProject accessions number PRJNA1129844. All other data and code used in this study are publicly and freely available through GitHub (https://github.com/Potnislab/pathogendiversity).

## Author contributions

NP conceptualized and designed the study. RB, AK, and IR processed the samples. RB, AK, and IR conducted analyses; specifically, RB and IR conducted ecological dynamics analysis. RB and IR, OR, and RM conducted statistical modeling analyses. AK worked on building pangenome and removal of blacklisted genes and the evolutionary analyses. DB conducted the greenhouse experiment. SS conducted analysis of pathogen fitness contributions towards epidemics using StrainRanking. RB, AK, and NP worked on data interpretation. APK, ZS, PR, IM, TT, BD, and ES helped collect samples and metadata from different tomato farms across four states. RB, AK, and NP wrote the manuscript with contributions from all the authors.

## Supporting Information

### Supplementary Text

**S1 Appendix. Climatic shifts and extremes explain variation in BLS epidemics across the Southeastern United States**

### Supplementary Figures

**S1 Fig**. A total of 8 fields consistently sampled over all three years displayed variable disease pressures across three years. Disease severity score is based on percent leaf area affected and estimated using the Horsfall-Barratt scale, which ranges from 1 to 12, with scale of 1 being no disease and 12 being 100% defoliation (Horsfall and Barratt 1945).

**S2 Fig.** (**A)** Disease severity ratings for different sampling time points across different states of Southeastern US. **(B)** the absolute abundance and **(C)** Relative abundance of *Xp* collected from different states during 2020, 2021, and 2022.

**S3 Fig.** Plots are comparing the samples based on the farm size (commercial vs small scale farms) **(A)** disease severity ratings; **(B)** Shannon diversity & **(C)** Number of Sequence clusters (SCs) or pathogen lineages present in the samples. The pathogen diversity was significantly higher in commercial fields compared to small-scale farms, although disease severity values did not significantly vary.

**S4 Fig.** Stacked bar plot depicting the co-occurrence of multiple *Xp* lineages, spatial and temporal variations, the introduction of new lineages, turnover, and dominance shifts in individual fields across various states during the mid and end of the season for the years 2020, 2021, and 2022

**S5 Fig.** Boxplots showing **(A)** disease severity ratings **& (B)** Average nucleotide diversity for farms having presence of less than and more than 2 Sequence clusters (pathogen lineages).

**S6 Fig. (A)** Correlation plot showing the relationship between Shannon diversity of *Xp* lineages and disease severity across all samples. The presence of more *Xp* lineages in the field correlates with higher BLS disease severity; **(B)** Boxplot showing the Shannon diversity across the samples with low and high disease severity conditions. High DS includes the samples with disease severity scale ratings of more than and equal to 5, whereas low DS includes the farms with disease severity scale ratings of less than 5.

**S7 Fig.** Evaluating the fitness contributions of individual vs mixed infections by strains belonging to different sequence clusters under greenhouse conditions. Plants were dip-inoculated with ∼1 X 10^6^ of cell suspensions for each treatment. **(A)** Boxplot presenting the disease severity ratings evaluated using Horsfall-Barratt scale at day 7 and 15 of post-inoculation. The mixed infections showed higher disease severity compared to single infections**. (B)** pairwise-comparisons of AUDPC values across different pairs of treatments. Differences in means of disease development i.e. AUDPC for different treatment pairs are shown. Mean values of disease development with a 95% confidence interval were contrasted between treatments. Difference in means of disease development between treatments with an estimate 0 have no difference in means of disease development among pair of strains. Negative and positive values represent statistically significant treatments based on Tukey’s HSD test.

**S8 Fig.** Using StrainRanking, **(A)** Comparisons of different SCs with fitness estimates and their confidence intervals under main effects were made. These estimates focused on SC3, SC4, and SC6 (which were more abundant in samples), and a group of SCs (SC1, SC2, SC5, SC7, and SC8) that were categorized as low-frequency (due to their lower abundance). The bottom plot illustrates the growth curves of these SCs, estimating their contributions to the progression of disease severity from mid and end of summer**. (B)** Pathogen-pathogen interactions, specifically between the two dominant SCs (SC3 and SC4), were analyzed based on their combined frequencies being either below or above 20%. Fitness estimates with confidence intervals were compared when the summed frequencies of SC3 and SC4 were taken into account. Different interaction scenarios were considered, such as SC3 > 20% & SC4 < 20%, SC4 > 20% & SC3 < 20%, and both SC3 & SC4 at equal frequencies, along with SC6 and the low-frequency SCs. The growth curves in the bottom plot highlight how these interactions contributed to the progression of disease during the season. **(C)** Pathogen-environment interactions were examined by applying the frequencies of SC3 or SC4 to different environmental variables, depending on whether each variable was above or below its mean at the sampling point. The analysis assumed weighted mean of genetic frequencies to calculate fitness levels as: *p* x mid-frequency + (1 - *p*) x end-frequency, where (*p* = 0.90).

**S9 Fig.** Using StrainRanking: **(A)** Comparisons of different SCs with fitness estimates and their confidence intervals under main effects were made. These estimates focused on SC3, SC4, and SC6 (which were more abundant in samples), and a group of SCs (SC1, SC2, SC5, SC7, and SC8) that were categorized as low-frequency (due to their lower abundance). The bottom plot illustrates the growth curves of these SCs, estimating their contributions to the progression of disease severity from mid and end of summer**. (B)** Pathogen-pathogen interactions, specifically between the two dominant SCs (SC3 and SC4), were analyzed based on their combined frequencies being either below or above 20%. Fitness estimates with confidence intervals were compared when the summed frequencies of SC3 and SC4 were taken into account. Different interaction scenarios were considered, such as SC3 > 20% & SC4 < 20%, SC4 > 20% & SC3 < 20%, and both SC3 & SC4 at equal frequencies, along with SC6 and the low-frequency SCs. The growth curves in the bottom plot highlight how these interactions contributed to the progression of disease during the season. **(C)** Pathogen-environment interactions were examined by applying the frequencies of SC3 or SC4 to different environmental variables, depending on whether each variable was above or below its mean at the sampling point. The analysis assumed weighted mean of genetic frequencies to calculate fitness levels as: *p* x mid-frequency + (1 - *p*) x end-frequency, where (*p* = 0.10).

**S10 Fig.** plot showing the average allele frequencies for those alleles having **(A)** *f >= 0.8* & **(B)** *f <0.8* during End-season 2020; **(C)** *f >= 0.8* & **(D)** *f <0.8* during Fall-season 2020 and traced for the frequency changes during the previous and following seasons of three years.

**S11 Fig.** Sankey plot shows counts of the alleles present in parallel across farms during **A)** End-Season 2020 & **B)** Fall-season of 2020 with frequency less than and equal to 0.2 and then counts for those alleles which stayed as major during the next seasons.

**S12 Fig.** In circos plot is presenting LH3 genome with all the colored bars indicating different genes in different scaffolds, where genes under positive selection are shown with the red colored asterisk sign in the grey colored ring and the red colored arrows are showing those genes which were present in parallel across farms and seasons.

### Supplementary Tables

**S1A Table.** Farm details and time of sampling during three sampling years

**S1B Table.** Details of farm and disease severity across season and sampling years used in the study

**S2 Table.** Climatic parameters used in the study and their meaning

**S3A Table.** Influence of various factors in BLS disease severity

**S3B Table.** Influence of various factors in absolute abundance of X. euvesicatoria pv. perforans

**S3C Table.** Influence of various factors in relative abundance of X. euvesicatoria pv. perforans

**S4 Table.** Coefficients Estimates and p-values for different Sequence Clusters

**S5 Table.** Gene under positive selection pressure (Annotations according LH3 Genome)

