## Supplementary file for "Interaction between climatic variation and pathogen diversity shape endemic disease dynamics in the agricultural settings"

Short title: Pathogen diversity in bacterial spot disease epidemics

**Supplementary Text:**

**S1 Appendix. Climatic shifts and extremes explain variation in BLS epidemics across the Southeastern United States**

We ran the logistic ordinal regression model to understand drivers for disease severity which indicated that the Shannon diversity of various lineages of *Xp* present in the field (t value = 4.15, $p < 0.05$) strongly influenced BLS disease severity as suggested in the earlier result (Fig. S5). Among the climatic factors, the standard deviation of Wet Bulb Temperature at 2 meters (t-value = 2.06, $p < 0.05$) and the standard deviation of the average of the wind direction at 10 meters above the surface of the earth (t-value = 2.73, $p < 0.05$) significantly increases BLS disease severity (Table 1A, Table S3A), indicating that more variations of wet bulb temperature (adiabetic saturation temperature) and wind direction values led to higher disease severity. In contrast, the skewness of clear sky surface photosynthetically active radiation (PAR) total (t-value = -2.57, $p <$ 0.05, negative skewness indicative of more data points accumulating in the upper range of clear sky PAR) and kurtosis of Temperature at 2 Meters Range (t-value = -2.12, $p < 0.05$, negative kurtosis indicative of when tail min temperature relatively close to typical minimum temperature) decrease the BLS disease severity (Table 1A, Table S3A). More specifically, for the skewness predictor, this implies that as the majority of clear sky PAR measurements accumulate towards the lower range (and hence have fewer extremes in this range), the less the BLS disease severity occurs; for the kurtosis predictor we also see that when the temperatures at 2 meters range have more extreme behaviors, the less is the BLS disease severity.

Next, a beta regression model predicting the drivers of *Xp* abundance identified one key predictor: the standard deviation of wind direction at 10 meters above the earth's surface. This variable has positive association with the absolute (z-value = 2.08, $p < 0.05$)) and relative (z-value = 2.69, $p <$

0.05) abundance of *Xp*, contributing to disease severity. In addition to this, the skewness of surface pressure (z $value_{abs}$ = -3.261, z $value_{rel}$ = -2.949, $p < 0.05$) and kurtosis of relative humidity at 2 meters (z $value_{abs}$ = -2.524, z $value_{rel}$ = -2.744, $p < 0.05$) showed a significant negative effect both on the absolute (Table 1B, Table S3B) and relative abundance of *Xp* (Table S3C). Thus, more variations in wind direction and extreme changes in surface pressure and relative humidity all appear to influence *Xp* abundance. Other significant predictors of *Xp* relative abundance included farm scale, with commercial farms having a larger abundance (z = 2.875, $p < 0.05$) (Table S3C). Finally, we used Dirichlet compositional regression to evaluate the influence of climatic factors in explaining the differential dynamics of pathogen lineages across different fields. This type of regression ensures that dependence between cluster abundance is accounted for (i..e. the proportion of clusters must sum to one). We found that the variables that appeared to be significant across pathogen lineages were the standard deviation of specific humidity at 2 meters and mid-season (compared to end-season). In particular, these appeared to diminish abundance across all clusters except for SC6. Also, kurtosis of relative humidity appears to have a negative impact on abundance in clusters SC1 to SC4 and SC8. In general, the contribution of the variables appears to be more significant in clusters SC3 and SC4 (and to a lesser extent SC2). The standard deviation of relative humidity appears to increase SC3 abundance. In contrast, variables standard deviation of minimum to the maximum temperature range at 2 meters, skewness of temperature at 2 meters, skewness of wind speed at 10 meters, kurtosis of relative humidity, and kurtosis of all-sky PAR appears to decrease the abundance of SC3. The variables, standard deviation of relative humidity, skewness of temperature at 2m, and Commercial scale farms (compared to small farms), appear to increase the abundance of SC4 (Table S4).

### Supplementary Figures:

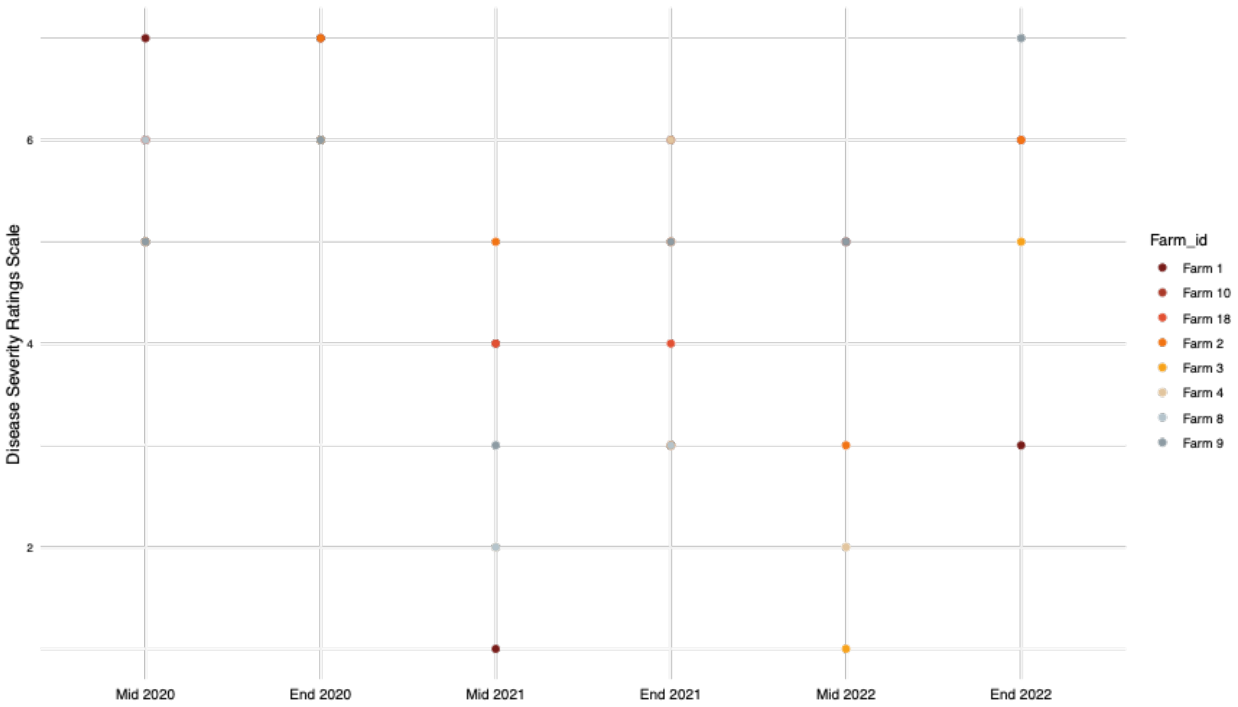


**S1 Fig**. A total of 8 fields consistently sampled over all three years displayed variable disease pressures across three years. Disease severity score is based on percent leaf area affected and estimated using the Horsfall-Barratt scale, which ranges from 1 to 12, with scale of 1 being no disease and 12 being 100% defoliation (Horsfall and Barratt 1945).

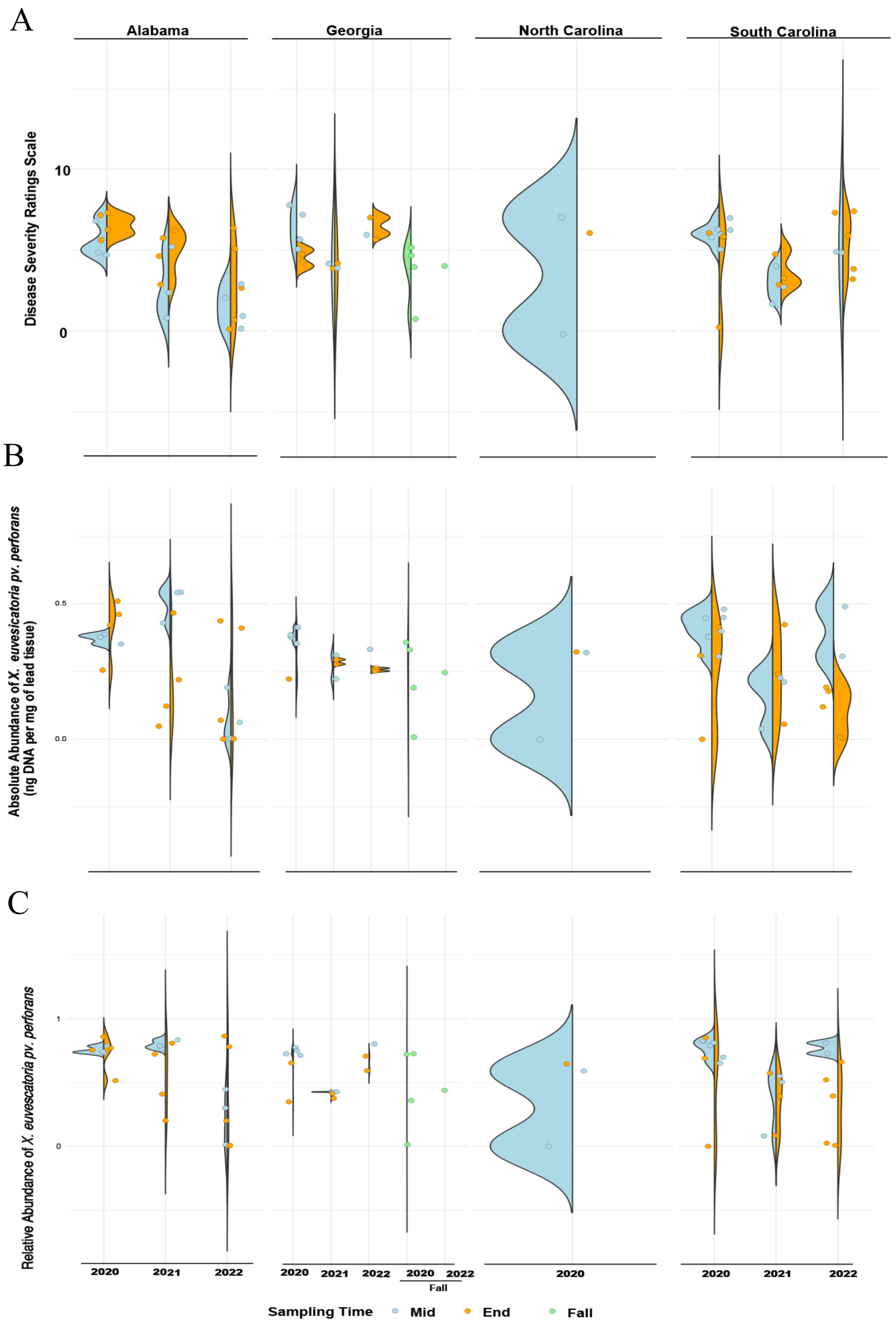

A
Alabama
Georgia
North Carolina
South Carolina
Disease Severity Ratings Scale
10
0
B
Absolute Abundance of X. euvesicatoria pv. perforans (ng DNA per mg of lead tissue)
0.5
0.0
C
Relative Abundance of X. euvesicatoria pv. perforans
1
0
2020
2021
2022
Fall
Sampling Time
Mid
End
Fall

**S2 Fig.** **(A)** Disease severity ratings for different sampling time points across different states of Southeastern US. **(B)** the absolute abundance and **(C)** Relative abundance of *Xp* collected from different states during 2020, 2021, and 2022.

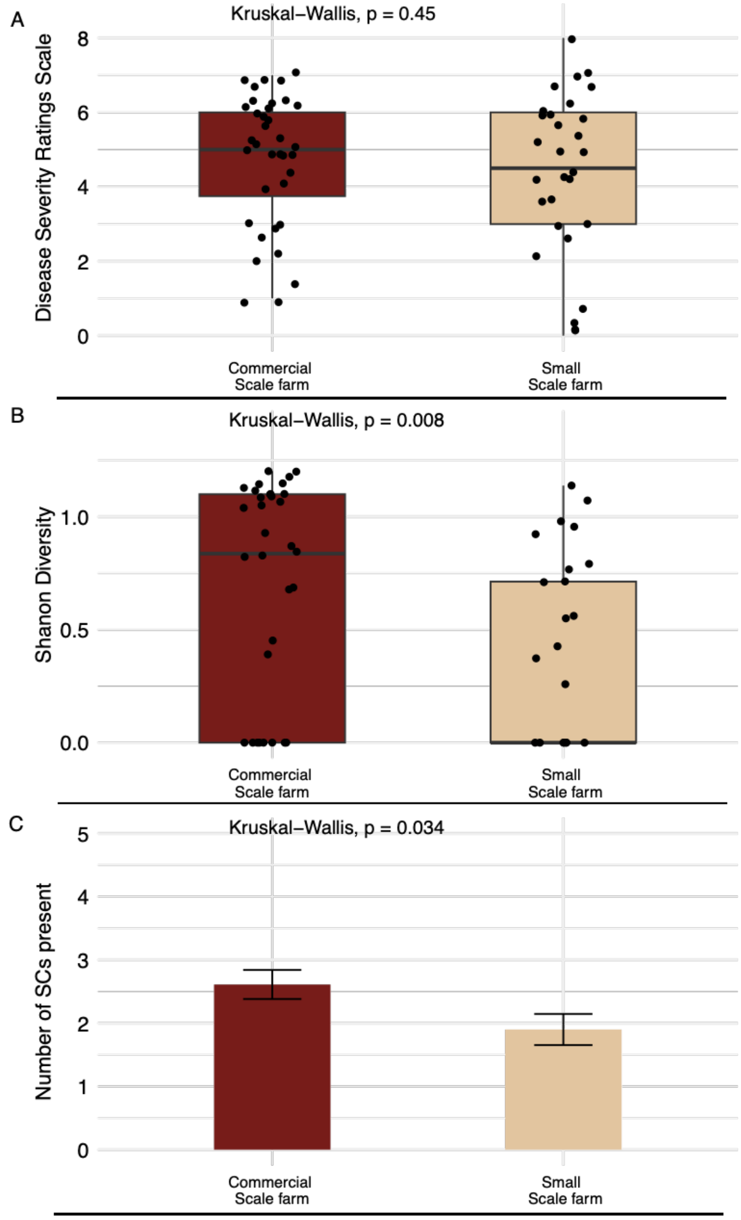


**S3 Fig.** Plots are comparing the samples based on the farm size (commercial vs small scale farms) **(A)** disease severity ratings; **(B)** Shannon diversity & **(C)** Number of Sequence clusters (SCs) or pathogen lineages present in the samples. The pathogen diversity was significantly higher in commercial fields compared to small-scale farms, although disease severity values did not significantly vary.

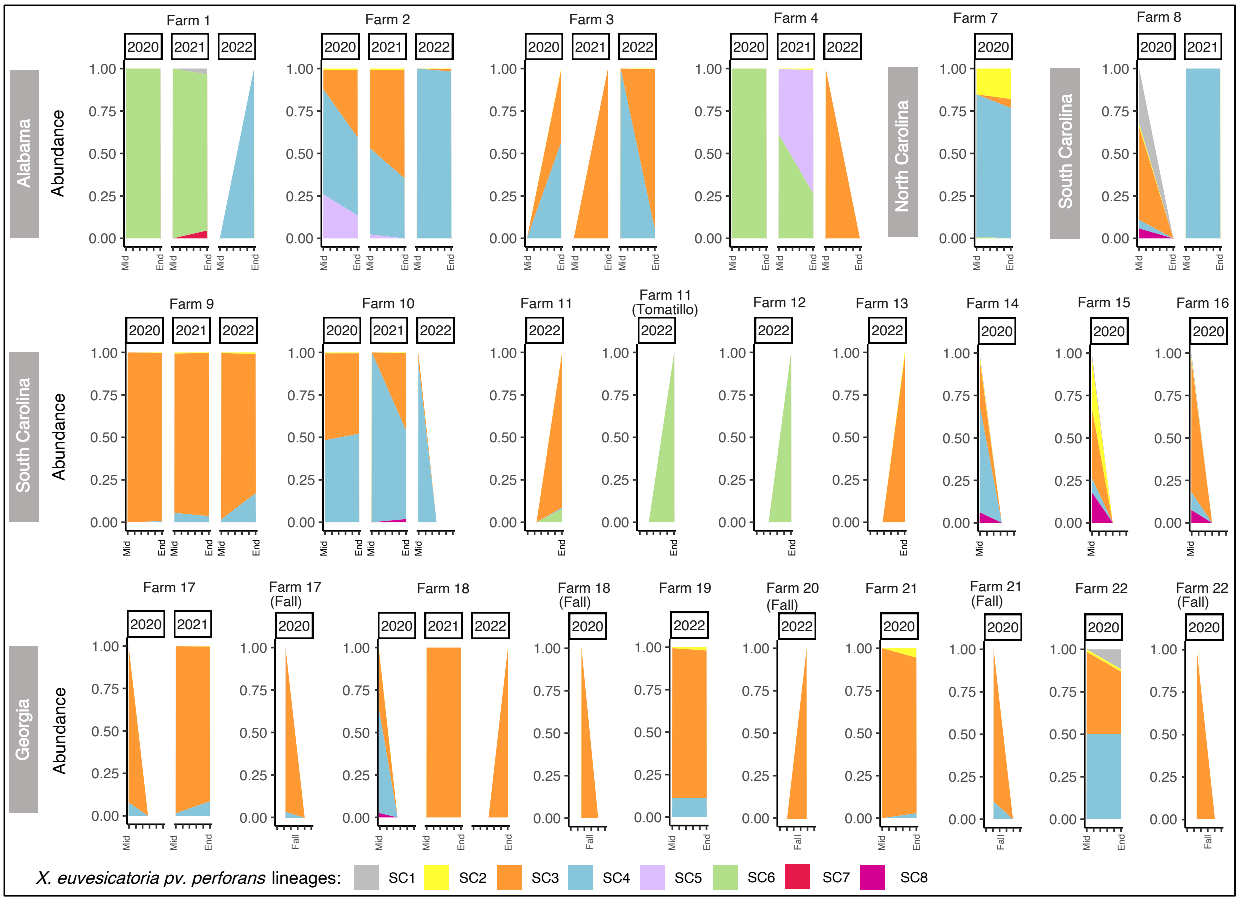


**S4 Fig.** Stacked bar plot depicting the co-occurrence of multiple *Xp* lineages, spatial and temporal variations, the introduction of new lineages, turnover, and dominance shifts in individual fields across various states during the mid and end of the season for the years 2020, 2021, and 2022

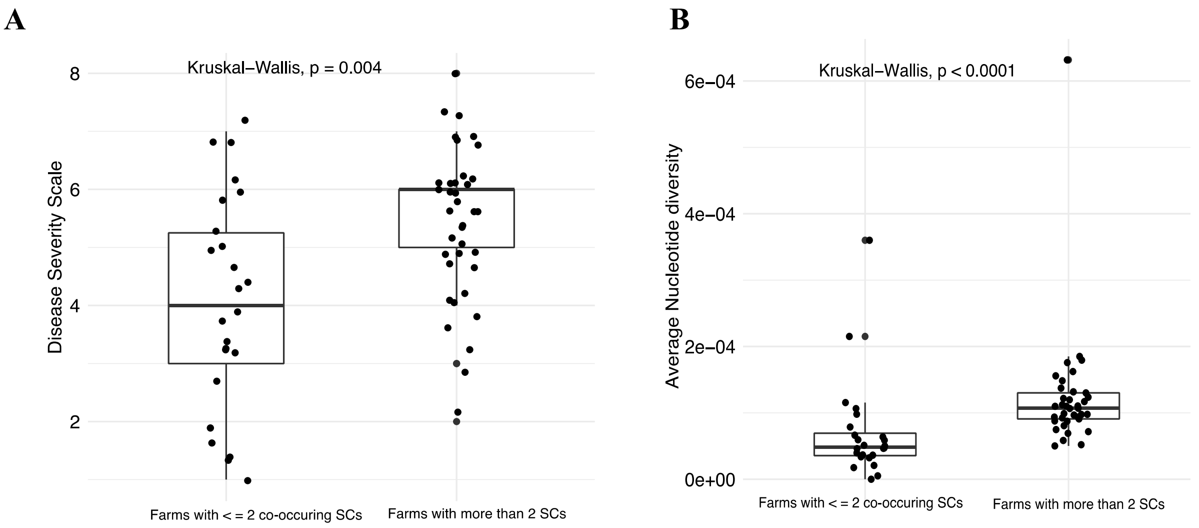


**S5 Fig.** Boxplots showing **(A)** disease severity ratings **& (B)** Average nucleotide diversity for farms having presence of less than and more than 2 Sequence clusters (pathogen lineages).

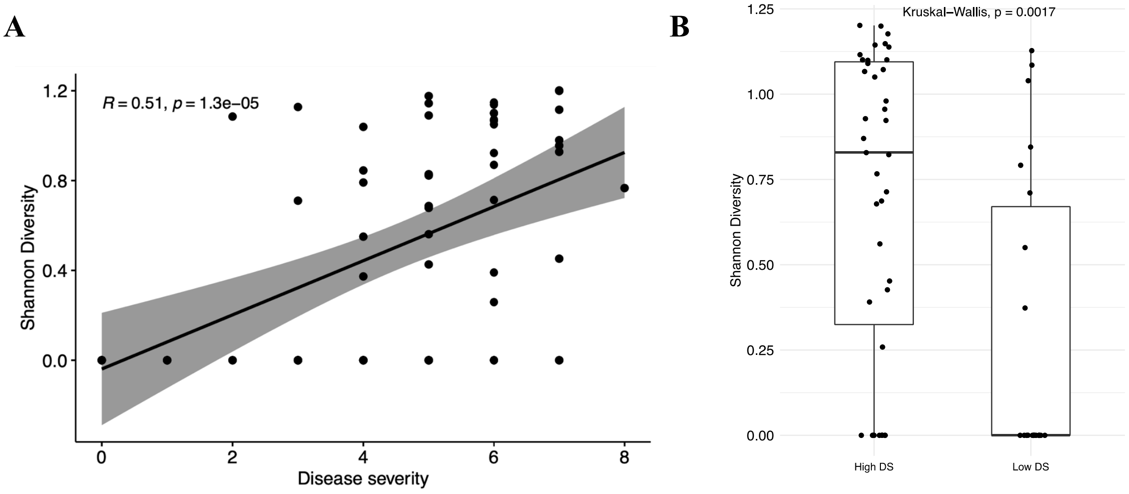


**S6 Fig. (A)** Correlation plot showing the relationship between Shannon diversity of *Xp* lineages and disease severity across all samples. The presence of more *Xp* lineages in the field correlates with higher BLS disease severity; **(B)** Boxplot showing the Shannon diversity across the samples with low and high disease severity conditions. High DS includes the samples with disease severity scale ratings of more than and equal to 5, whereas low DS includes the farms with disease severity scale ratings of less than 5.

A

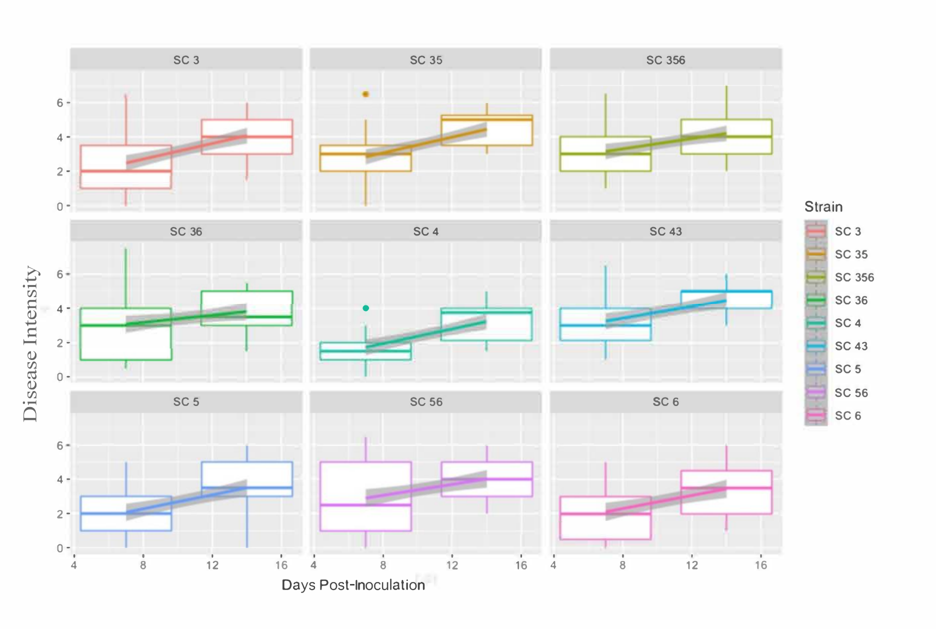
SC 3
SC 35
SC 356
SC 36
SC 4
SC 43
SC 5
SC 56
SC 6
Disease Intensity
Days Post-Inoculation
Strain

B

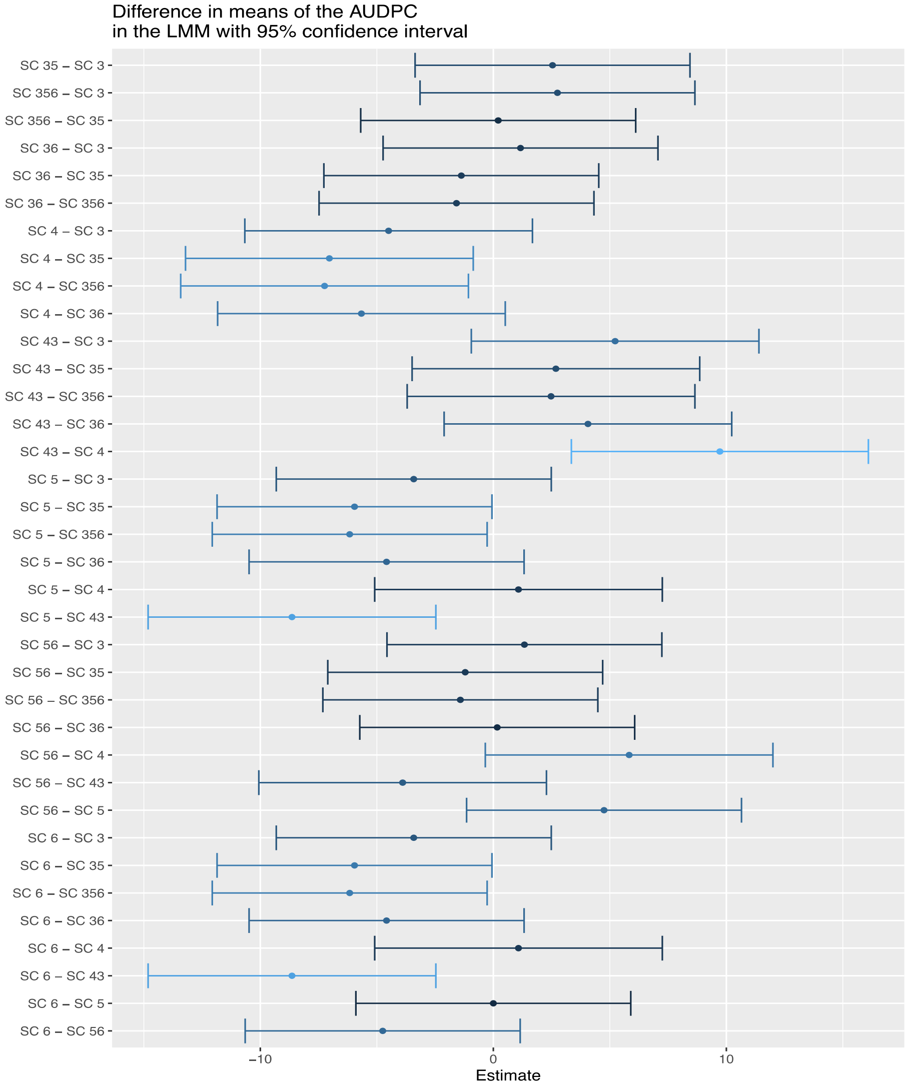
Difference in means of the AUDPC
in the LMM with 95% confidence interval
SC 35 – SC 3
SC 356 – SC 3
SC 356 – SC 35
SC 36 – SC 3
SC 36 – SC 35
SC 36 – SC 356
SC 4 – SC 3
SC 4 – SC 35
SC 4 – SC 356
SC 4 – SC 36
SC 43 – SC 3
SC 43 – SC 35
SC 43 – SC 356
SC 43 – SC 36
SC 43 – SC 4
SC 5 – SC 3
SC 5 – SC 35
SC 5 – SC 356
SC 5 – SC 36
SC 5 – SC 4
SC 5 – SC 43
SC 56 – SC 3
SC 56 – SC 35
SC 56 – SC 356
SC 56 – SC 36
SC 56 – SC 4
SC 56 – SC 43
SC 56 – SC 5
SC 6 – SC 3
SC 6 – SC 35
SC 6 – SC 356
SC 6 – SC 36
SC 6 – SC 4
SC 6 – SC 43
SC 6 – SC 5
SC 6 – SC 56
Estimate

**S7 Fig.** Evaluating the fitness contributions of individual vs mixed infections by strains belonging to different sequence clusters under greenhouse conditions. Plants were dip-inoculated with ~1 X $10^6$ of cell suspensions for each treatment. **(A)** Boxplot presenting the disease severity ratings evaluated using Horsfall-Barratt scale at day 7 and 15 of post-inoculation. The mixed infections showed higher disease severity compared to single infections**. (B)** pairwise-comparisons of AUDPC values across different pairs of treatments. Differences in means of disease development i.e. AUDPC for different treatment pairs are shown. Mean values of disease development with a 95% confidence interval were contrasted between treatments. Difference in means of disease development between treatments with an estimate 0 have no difference in means of disease development among pair of strains. Negative and positive values represent statistically significant treatments based on Tukey's HSD test.

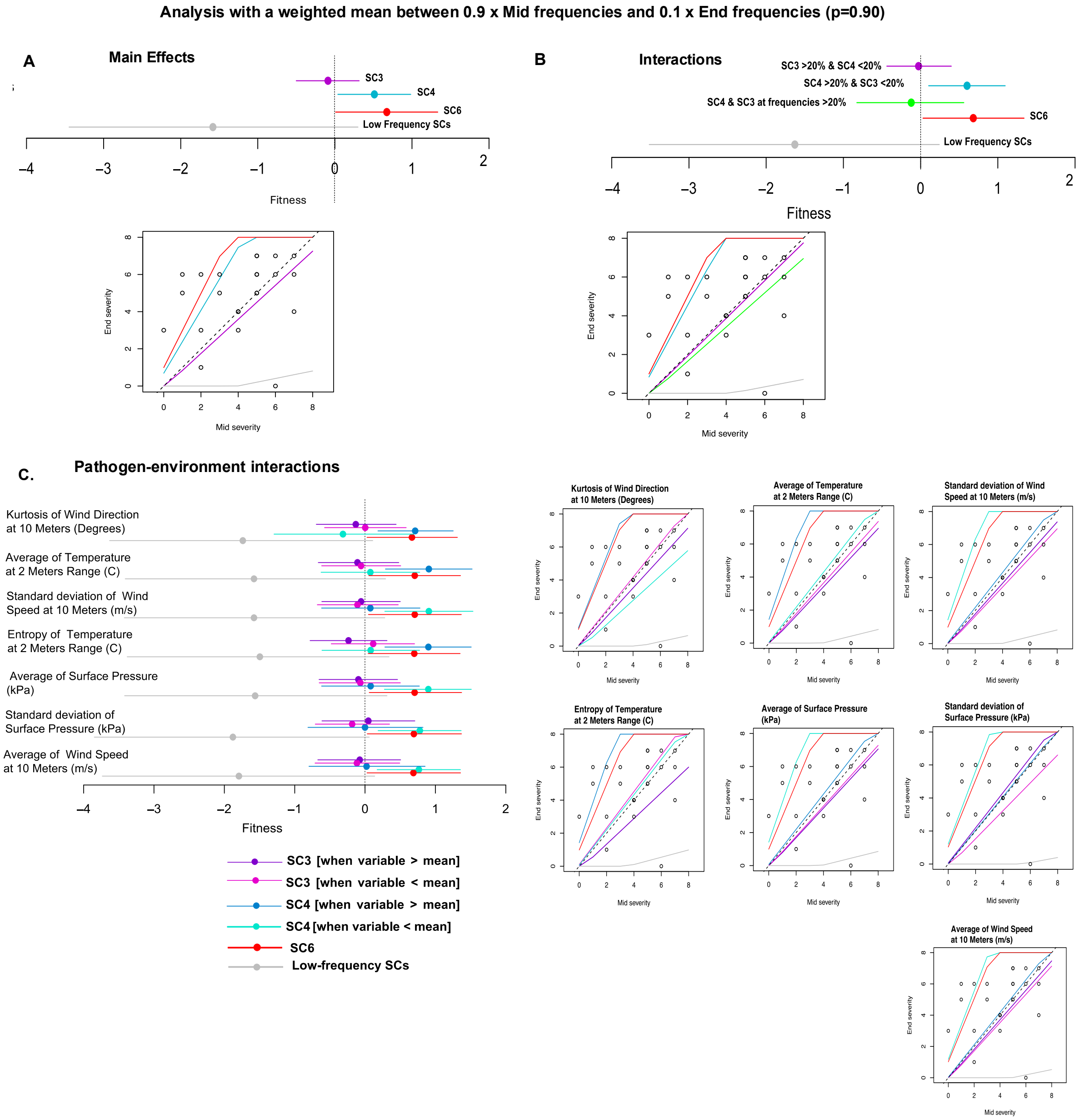


**S8 Fig.** Using StrainRanking, **(A)** Comparisons of different SCs with fitness estimates and their confidence intervals under main effects were made. These estimates focused on SC3, SC4, and SC6 (which were more abundant in samples), and a group of SCs (SC1, SC2, SC5, SC7, and SC8) that were categorized as low-frequency (due to their lower abundance). The bottom plot illustrates the growth curves of these SCs, estimating their contributions to the progression of disease severity from mid and end of summer**. (B)** Pathogen-pathogen interactions, specifically between the two dominant SCs (SC3 and SC4), were analyzed based on their combined frequencies being either below or above 20%. Fitness estimates with confidence intervals were compared when the summed frequencies of SC3 and SC4 were taken into account. Different interaction scenarios were considered, such as SC3 > 20% & SC4 < 20%, SC4 > 20% & SC3 < 20%, and both SC3 & SC4 at equal frequencies, along with SC6 and the low-frequency SCs. The growth curves in the bottom plot highlight how these interactions contributed to the progression of disease during the season. **(C)** Pathogen-environment interactions were examined by applying the frequencies of SC3 or SC4 to different environmental variables, depending on whether each variable was above or below its mean at the sampling point. The analysis assumed weighted mean of genetic frequencies to calculate fitness levels as: $p$ x mid-frequency + (1 - $p$) x end-frequency, where ($p$ = 0.90).

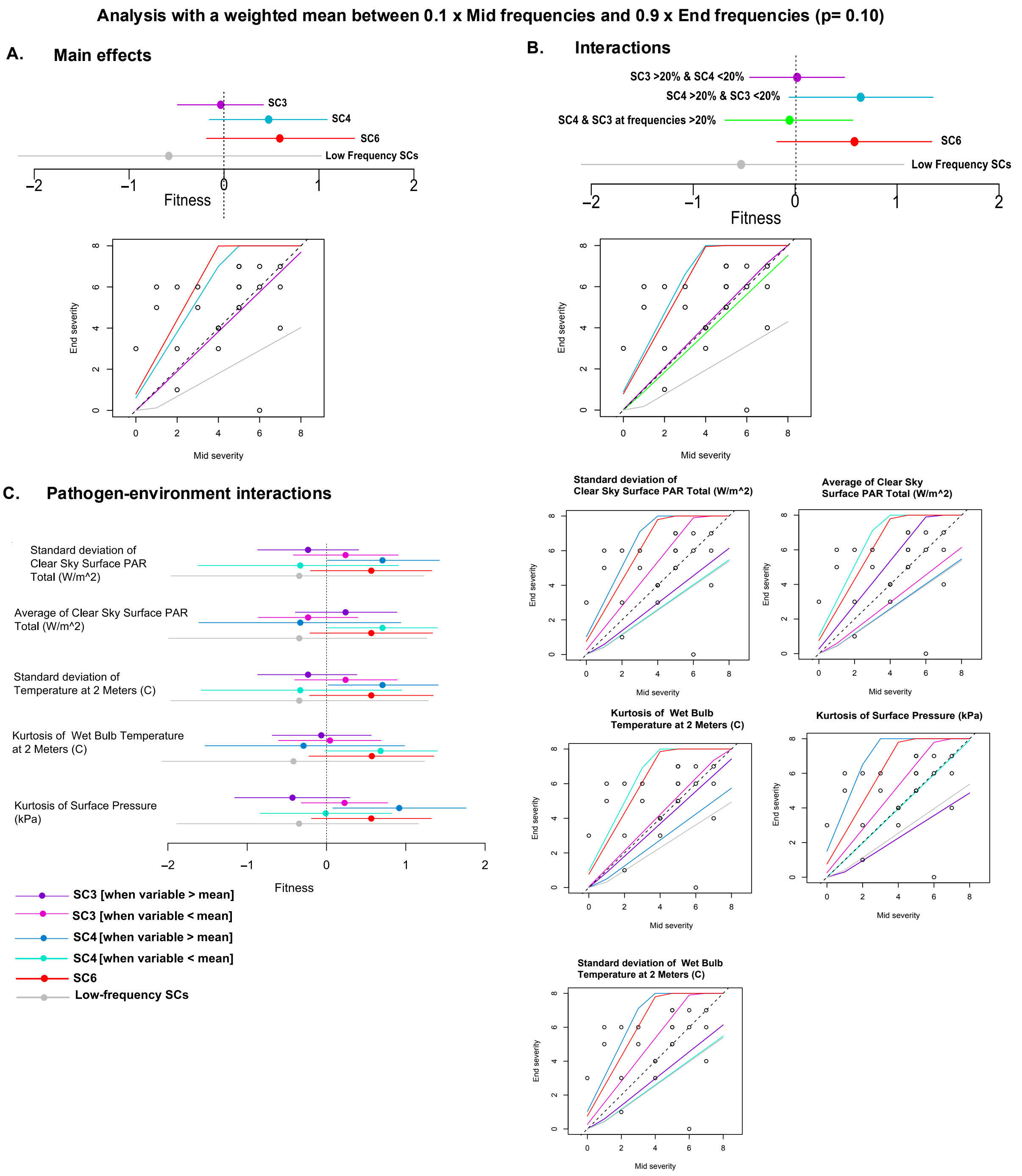


**S9 Fig.** Using StrainRanking: **(A)** Comparisons of different SCs with fitness estimates and their confidence intervals under main effects were made. These estimates focused on SC3, SC4, and SC6 (which were more abundant in samples), and a group of SCs (SC1, SC2, SC5, SC7, and SC8) that were categorized as low-frequency (due to their lower abundance). The bottom plot illustrates the growth curves of these SCs, estimating their contributions to the progression of disease severity from mid and end of summer**. (B)** Pathogen-pathogen interactions, specifically between the two dominant SCs (SC3 and SC4), were analyzed based on their combined frequencies being either below or above 20%. Fitness estimates with confidence intervals were compared when the summed frequencies of SC3 and SC4 were taken into account. Different interaction scenarios were considered, such as SC3 > 20% & SC4 < 20%, SC4 > 20% & SC3 < 20%, and both SC3 & SC4 at equal frequencies, along with SC6 and the low-frequency SCs. The growth curves in the bottom plot highlight how these interactions contributed to the progression of disease during the season. **(C)** Pathogen-environment interactions were examined by applying the frequencies of SC3 or SC4 to different environmental variables, depending on whether each variable was above or below its mean at the sampling point. The analysis assumed weighted mean of genetic frequencies to calculate fitness levels as: $p$ x mid-frequency + (1 - $p$) x end-frequency, where ($p$ = 0.10).

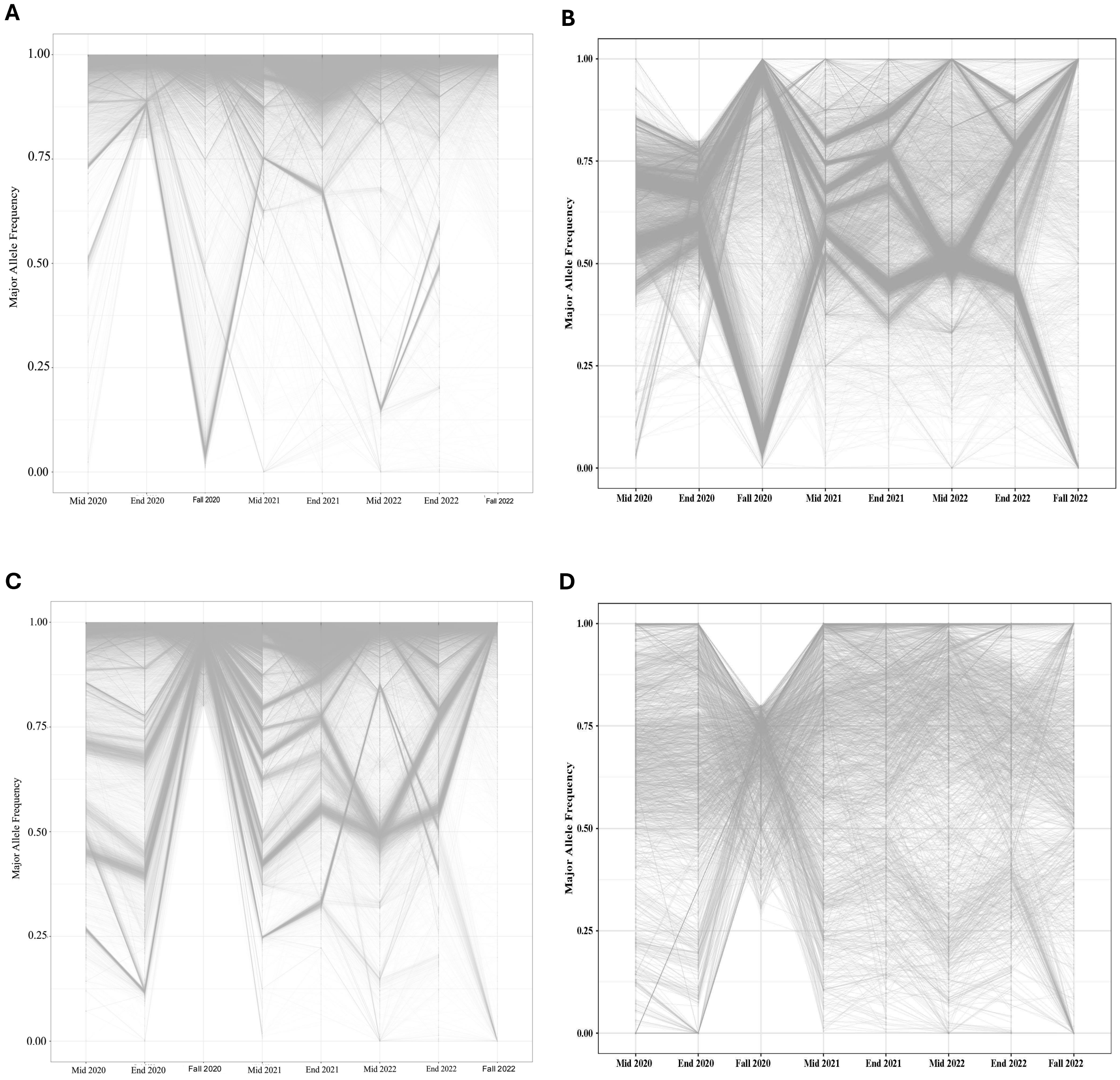


**S10 Fig.** plot showing the average allele frequencies for those alleles having **(A)** *f >= 0.8* & **(B)** *f <0.8* during End-season 2020; **(C)** *f >= 0.8* & **(D)** *f <0.8* during Fall-season 2020 and traced for the frequency changes during the previous and following seasons of three years.

**A**

**B**

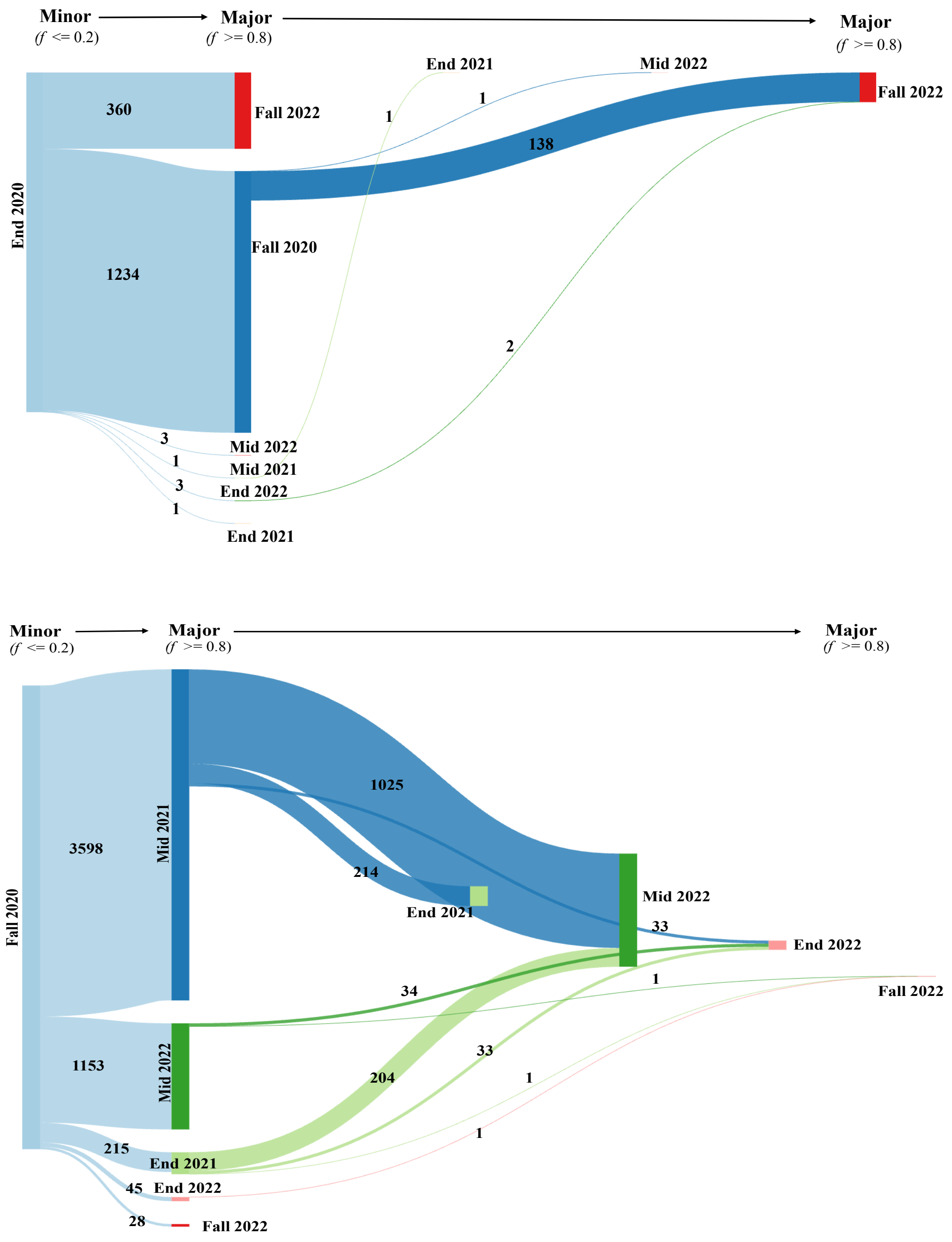


**S11 Fig.** Sankey plot shows counts of the alleles present in parallel across farms during **A)** End-Season 2020 & **B)** Fall-season of 2020 with frequency less than and equal to 0.2 and then counts for those alleles which stayed as major during the next seasons.

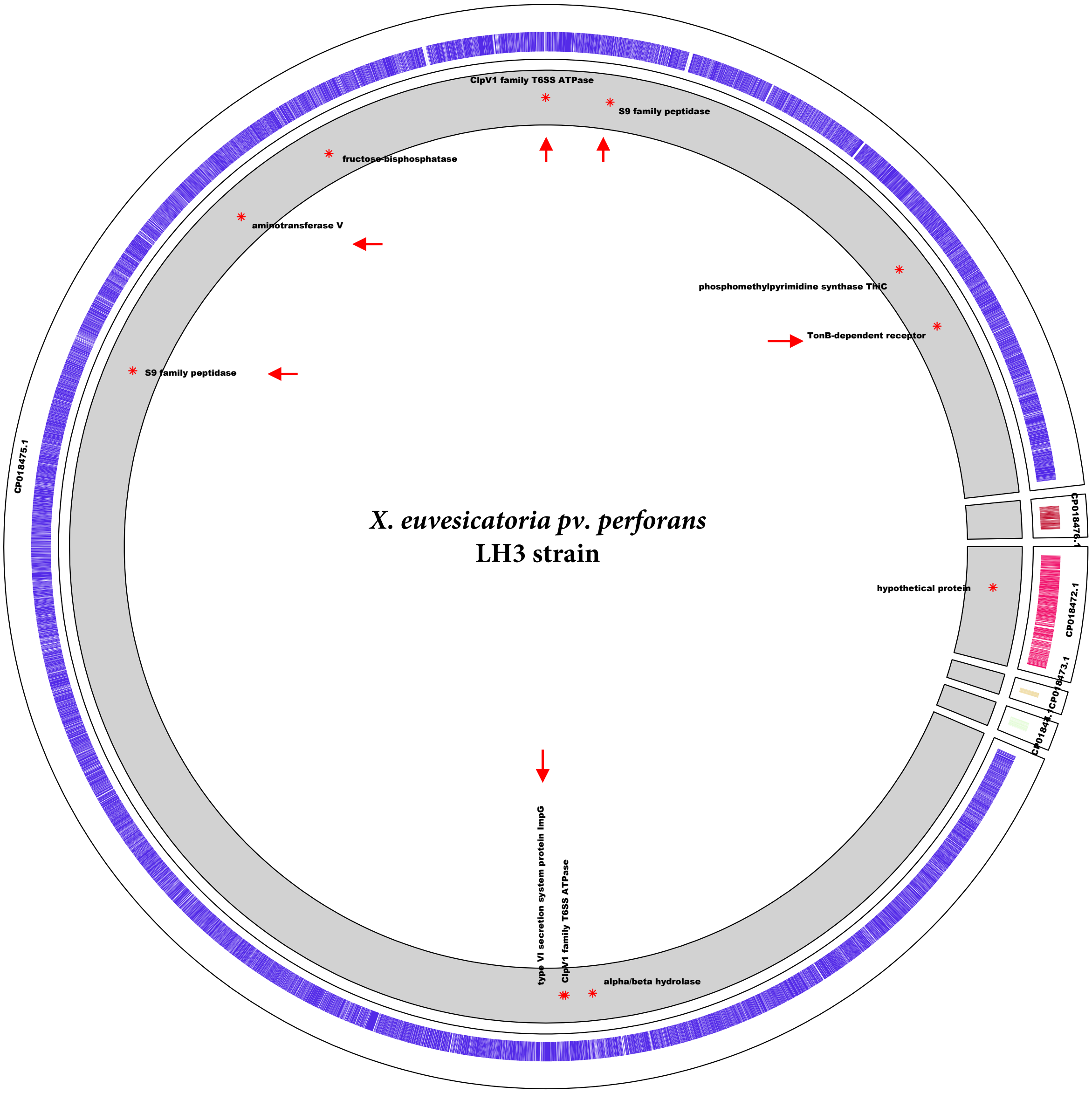


**S12 Fig.** In circos plot is presenting LH3 genome with all the colored bars indicating different genes in different scaffolds, where genes under positive selection are shown with the red colored asterisk sign in the grey colored ring and the red colored arrows are showing those genes which were present in parallel across farms and seasons.

**Supplementary Tables:**

| S1A Table. Farm details and time of sampling during three sampling years | | | | | | | |
|---|---|---|---|---|---|---|---|
| **Farm ID** | **Year** | **Farm scale** | **State** | **Growing Season Start Date** | **Mid Season Sampling Date (Summer)** | **End Season Sampling Date (Summer)** | **Fall Season Sampling Date** |
| Farm 1 | 2020 | Small | Alabama | 4/6/20 | 5/21/20 | 7/17/20 | |
| Farm 1 | 2021 | Small | Alabama | 4/12/21 | 5/27/21 | 6/17/21 | |
| Farm 1 | 2022 | Small | Alabama | 4/13/22 | 5/28/22 | 6/30/22 | |
| Farm 2 | 2020 | Commercial | Alabama | 4/6/20 | 5/21/20 | 6/17/20 | |
| Farm 2 | 2021 | Commercial | Alabama | 4/12/21 | 5/27/21 | 6/17/21 | |
| Farm 2 | 2022 | Commercial | Alabama | 4/13/22 | 5/28/22 | 6/30/22 | |
| Farm 3 | 2020 | Commercial | Alabama | 5/19/20 | 7/3/20 | 7/28/20 | |
| Farm 3 | 2021 | Commercial | Alabama | 5/24/21 | 7/8/21 | 7/17/21 | |
| Farm 3 | 2022 | Commercial | Alabama | 5/27/22 | 7/11/22 | 8/17/22 | |
| Farm 4 | 2020 | Commercial | Alabama | 4/13/20 | 5/28/20 | 7/3/20 | |
| Farm 4 | 2021 | Commercial | Alabama | 4/17/21 | 6/1/21 | 7/1/21 | |
| Farm 4 | 2022 | Commercial | Alabama | 4/9/22 | 5/24/22 | 6/16/22 | |
| Farm 5 | 2022 | Small | Alabama | 5/12/22 | | 8/10/22 | |
| Farm 6 | 2020 | Small | North Carolina | 5/31/20 | 7/15/20 | 7/30/20 | |
| Farm 7 | 2020 | Small | North Carolina | 5/31/20 | 7/15/20 | 8/3/20 | |
| Farm 8 | 2020 | Small | South Carolina | 4/5/20 | 5/20/20 | 6/20/20 | |
| Farm 8 | 2021 | Small | South Carolina | 4/6/21 | 5/21/21 | 6/21/21 | |
| Farm 9 | 2020 | Small | South Carolina | 4/4/20 | 5/19/20 | 6/17/20 | |
| Farm 9 | 2021 | Small | South Carolina | 4/6/21 | 5/21/21 | 6/24/21 | |
| Farm 9 | 2022 | Small | South Carolina | 4/5/22 | 5/20/22 | 6/14/22 | |
| Farm 10 | 2020 | Commercial | South Carolina | 4/4/20 | 5/19/20 | 6/17/20 | |
| Farm 10 | 2021 | Commercial | South Carolina | 5/3/21 | 6/17/21 | 7/17/21 | |
| Farm 10 | 2022 | Commercial | South Carolina | 4/5/22 | 5/20/22 | 6/18/22 | |
| Farm 11-1 | 2022 | Commercial | South Carolina | 5/17/22 | | 8/16/22 | |
| Farm 11-2 | 2022 | Commercial | South Carolina | 5/17/22 | | 8/16/22 | |
| Farm 12 | 2022 | Commercial | South Carolina | 5/17/22 | | 8/16/22 | |
| Farm 13 | 2022 | Commercial | South Carolina | 5/17/22 | | 8/16/22 | |
| Farm 14 | 2020 | Commercial | South Carolina | 6/7/20 | 7/22/20 | | |
| Farm 15 | 2020 | Commercial | South Carolina | 6/7/20 | 7/22/20 | | |
| Farm 16 | 2020 | Commercial | South Carolina | 6/7/20 | 7/22/20 | | |

| | | | | | | | |
|---|---|---|---|---|---|---|---|
| Farm 17 | 2020 | Small | Georgia | 5/3/20 | 6/17/20 | | |
| Farm 17 | 2021 | Small | Georgia | 5/3/21 | 6/17/21 | 7/17/21 | |
| Farm 17-W | 2020 | Small | Georgia | 8/15/21 | | | 9/30/20 |
| Farm 18 | 2020 | Small | Georgia | 5/3/20 | 6/17/20 | | |
| Farm 18-W | 2020 | Small | Georgia | 8/15/20 | | | 9/30/20 |
| Farm 18 | 2021 | Small | Georgia | 5/3/21 | 6/17/21 | 7/17/21 | |
| Farm 18 | 2022 | Small | Georgia | 5/4/22 | | 7/6/22 | |
| Farm 19 | 2022 | Commercial | Georgia | 5/1/22 | 6/15/22 | 7/7/22 | |
| Farm 20-W | 2022 | Small | Georgia | 8/15/22 | | | 10/6/22 |
| Farm 21 | 2020 | Commercial | Georgia | 5/12/20 | 6/26/20 | 7/26/20 | |
| Farm 21-W | 2020 | Commercial | Georgia | 8/15/20 | | | 9/30/20 |
| Farm 22 | 2020 | Commercial | Georgia | 5/12/20 | 6/26/20 | 7/26/20 | |
| Farm 22-W | 2020 | Commercial | Georgia | 8/15/20 | | | 9/30/20 |

**S1B Table. Details of farm and disease severity across season and sampling years used in the study**

| **Farm ID** | **State** | **Sampling time** | **Sample ID (2020)** | **Average Disease Severity (2020)** | **Sample ID (2021)** | **Average Disease Severity (2021)** | **Sample ID (2022)** | **Average Disease Severity (2022)** |
|---|---|---|---|---|---|---|---|---|
| Farm 1 | Alabama | Mid-season | B_ALM | 7 | BALM | 1 | BALM22 | 0 |
| Farm 1 | Alabama | End-season | B_ALE | 7 | BALE | 6 | BALE22 | 3 |
| Farm 2 | Alabama | Mid-season | A_ALM | 5 | AALM | 5 | AALM22 | 3 |
| Farm 2 | Alabama | End-season | A_ALE | 7 | AALE | 5 | AALE22 | 6 |
| Farm 3 | Alabama | Mid-season | L_ALM | 2 | LALM | 1 | LALM22 | 1 |
| Farm 3 | Alabama | End-season | L_ALE | 6 | LALE | 3 | LALE22 | 5 |
| Farm 4 | Alabama | Mid-season | S_ALM | 5 | SALM | 2 | SALM22 | 2 |
| Farm 4 | Alabama | End-season | S_ALE | 6 | SALE | 6 | SALE22 | 0 |
| Farm 5 | Alabama | End-season | | | | | EVS22 | 0 |

| | | | | | | | | |
|---|---|---|---|---|---|---|---|---|
| Farm 6 | North Carolina | Mid-season | F_NCM | 0 | | | | |
| Farm 7 | North Carolina | Mid-season | L_NCM | 7 | | | | |
| Farm 7 | North Carolina | End-season | L_NCE | 6 | | | | |
| Farm 8 | South Carolina | Mid-season | U_SCM | 6 | BSCM | 2 | | |
| Farm 8 | South Carolina | End-season | U_SCE | 0 | BSCE | 3 | | |
| Farm 9 | South Carolina | Mid-season | D_SCM | 5 | DSCM | 3 | DSCM22 | 5 |
| Farm 9 | South Carolina | End-season | D_SCE | 6 | DSCE | 5 | DSCE22 | 7 |
| Farm 10 | South Carolina | Mid-season | S_SCM | 6 | SSCM | 4 | SSCM22 | 5 |
| Farm 10 | South Carolina | End-season | S_SCE | 6 | SSCE | 3 | | |
| Farm 11-1 | South Carolina | End-season | | | | | PUSC22 | 4 |
| Farm 11-2 | South Carolina | End-season | | | | | STSC22 | 3 |
| Farm 12 | South Carolina | End-season | | | | | ESC22 | 6 |
| Farm 13 | South Carolina | End-season | | | | | WSC22 | 7 |
| Farm 14 | South Carolina | Mid-season | D1_SCM | 6 | | | | |
| Farm 15 | South Carolina | Mid-season | D2_SCM | 7 | | | | |
| Farm 16 | South Carolina | Mid-season | D3_SCM | 6 | | | | |
| Farm 17 | Georgia | Mid-season | C_GAM | 8 | CGAM | 4 | | |
| Farm 17 | Georgia | End-season | | | CGAE | 4 | | |
| Farm 17-W | Georgia | Winter | F1_GAW | 4 | | | | |
| Farm 18 | Georgia | Mid-season | H_GAM | 6 | HGAM | 4 | | |
| Farm 18 | Georgia | End-season | | | HGAE | 4 | CHGA22 | 6 |
| Farm 18-W | Georgia | Fall | F3_GAW | 5 | | | | |
| Farm 19 | Georgia | Mid-season | | | | | NGAM22 | 6 |

| | | | | | | | | |
|---|---|---|---|---|---|---|---|---|
| Farm 19 | Georgia | End-season | | | | | NGAE22 | 7 |
| Farm 20-W | Georgia | Fall | | | | | RPGA22 | 4 |
| Farm 21 | Georgia | Mid-season | S_GAM | 5 | | | | |
| Farm 21 | Georgia | End-season | C_GAW | 5 | | | | |
| Farm 21-W | Georgia | Fall | F2_GAW | 5 | | | | |
| Farm 22 | Georgia | Mid-season | W_GAM | 7 | | | | |
| Farm 22 | Georgia | End-season | M_GAW | 4 | | | | |
| Farm 22-W | Georgia | Fall | F4_GAW | 1 | | | | |

| **S2 Table. Climatic parameters used in the study and their meaning** | |
|---|---|
| **Abbreviation** | **Parameters** |
| T2M | Temperature at 2 Meters (C) |
| T2MDEW | Dew/Frost Point at 2 Meters (C) |
| T2MWET | Wet Bulb Temperature at 2 Meters (C) |
| TS | Earth Skin Temperature (C) |
| T2M_RANGE | Temperature at 2 Meters Range (C) |
| QV2M | Specific Humidity at 2 Meters (g/kg) |
| RH2M | Relative Humidity at 2 Meters (%) |
| PRECTOTCORR | Precipitation Corrected (mm/day) |
| CLRSKY_SFC_PAR_TOT | Clear Sky Surface PAR Total (W/m^2) |
| ALLSKY_SFC_PAR_TOT | All Sky Surface PAR Total (W/m^2) |
| PS | Surface Pressure (kPa) |
| WS10M | Wind Speed at 10 Meters (m/s) |
| WD10M | Wind Direction at 10 Meters (Degrees) |
| ALLSKY_SFC_UV_INDEX | All Sky Surface UV Index (dimensi |
| ALLSKY_SFC_LW_DWN | All Sky Surface Longwave Downward |

| **S3A Table. Influence of various factors in BLS disease severity** | | | |
|---|---|---|---|
| **Factors** | **Value** | **Std. Error** | **t value** |
| Shannon diversity | 2.37409 | 0.57175 | 4.1523 |
| Average of Relative humidity at 2 Meters (%) | -0.09859 | 0.0819 | -1.2037 |

| | | | |
|---|---|---|---|
| Standard deviation of Wet Bulb Temperature at 2 Meters | 0.96797 | 0.46843 | 2.0664 |
| Standard deviation of Wind Direction at 10 Meters | 0.09547 | 0.03492 | 2.7338 |
| Skewness of Clear Sky Surface PAR Total | -0.94701 | 0.36778 | -2.5749 |
| Skewness of Surface Pressure | -0.19601 | 1.19397 | -0.1642 |
| Skewness of Wind Direction at 10 Meters | -1.8786 | 1.06701 | -1.7606 |
| Kurtosis of Temperature at 2 Meters Range (C) | -1.25559 | 0.59055 | -2.1262 |
| Kurtosis of All Sky Surface PAR Total | 0.29636 | 0.28523 | 1.039 |
| Entropy of Precipitation Corrected (mm/day) | 0.94536 | 0.52329 | 1.8066 |

**S3B Table. Influence of various factors in absolute abundance of X. euvesicatoria pv. perforans**

| Factors | Estimate | Std. Error | z value | Pr(>\|z\|) | Remark |
|---|---|---|---|---|---|
| (Intercept) | -2.3559 | 1.41676 | -1.663 | 0.09633 | . |
| Standard deviation of Wind Direction at 10 Meters | 0.03934 | 0.01885 | 2.087 | 0.03686 | * |
| Skewness of Relative humidity at 2 Meters (%) | 0.40492 | 0.35996 | 1.125 | 0.26063 | |
| Skewness of Surface Pressure | -1.70069 | 0.52146 | -3.261 | 0.00111 | ** |
| Skewness of Wind Speed at 10 Meters | 0.18791 | 0.30416 | 0.618 | 0.53672 | |
| Kurtosis of Relative humidity at 2 Meters (%) | -0.66357 | 0.26293 | -2.524 | 0.01161 | * |
| Sampling Time: Mid | 0.19688 | 0.26209 | 0.751 | 0.45254 | |
| Year: 2020 | -0.12068 | 0.28662 | -0.421 | 0.67372 | |
| Year: 2022 | -0.24572 | 0.33542 | -0.733 | 0.46382 | |

**S3C Table. Influence of various factors in relative abundance of X. euvesicatoria pv. perforans**

| Factors | Estimate | Std. Error | z value | Pr(>\|z\|) | Remarks |
|---|---|---|---|---|---|
| (Intercept) | -3.130515 | 3.651502 | -0.857 | 0.39127 | |
| Average of Wind Direction at 10 Meters | 0.006655 | 0.013484 | 0.494 | 0.62163 | |
| Standard deviation of Wind Direction at 10 Meters | 0.056105 | 0.020784 | 2.699 | 0.00695 | ** |
| Skewness of Surface Pressure | -1.839924 | 0.623977 | -2.949 | 0.00319 | ** |
| Kurtosis of Relative humidity at 2 Meters (%) | -0.716617 | 0.261137 | -2.744 | 0.00607 | ** |
| Kurtosis of Surface pressure | -0.30025 | 0.349703 | -0.859 | 0.39057 | |
| Sampling Time: Mid | 0.184459 | 0.295216 | 0.625 | 0.53208 | |
| Year: 2020 | -0.135931 | 0.299429 | -0.454 | 0.64985 | |

<table>
<tr><td>Production Scale: Commercial</td><td>0.753191</td><td>0.262011</td><td>2.875</td><td>0.00404</td><td>**</td></tr>
</table>

<table>
<tr><th colspan="6">S4 Table. Coefficients Estimates and p-values for different Sequence Clusters</th></tr>
<tr><th colspan="6">S4A Table. Coefficients Estimates and p-values for SC1</th></tr>
<tr><td></td><td>Estimate</td><td>Std. Error</td><td>z value</td><td>Pr(>|z|)</td><td>Remarks</td></tr>
<tr><td>(Intercept)</td><td>7.78989</td><td>10.17134</td><td>0.766</td><td>0.4438</td><td></td></tr>
<tr><td>Rel</td><td>2.30138</td><td>1.51712</td><td>1.517</td><td>0.1293</td><td></td></tr>
<tr><td>Abs</td><td>-3.51279</td><td>2.67922</td><td>-1.311</td><td>0.1898</td><td></td></tr>
<tr><td>Av_CLRSKY_SFC_PAR_TOT</td><td>-0.02553</td><td>0.06865</td><td>-0.372</td><td>0.71</td><td></td></tr>
<tr><td>Sd_T2M_RANGE</td><td>-0.49538</td><td>0.50816</td><td>-0.975</td><td>0.3296</td><td></td></tr>
<tr><td>Sd_QV2M</td><td>-1.16982</td><td>0.46306</td><td>-2.526</td><td>0.0115</td><td>*</td></tr>
<tr><td>Sd_RH2M</td><td>0.26875</td><td>0.16442</td><td>1.635</td><td>0.1022</td><td></td></tr>
<tr><td>Skew_T2M</td><td>-0.53049</td><td>0.53656</td><td>-0.989</td><td>0.3228</td><td></td></tr>
<tr><td>Skew_WS10M</td><td>0.16046</td><td>0.4179</td><td>0.384</td><td>0.701</td><td></td></tr>
<tr><td>Skew_WD10M</td><td>0.61428</td><td>0.70207</td><td>0.875</td><td>0.3816</td><td></td></tr>
<tr><td>Kur_RH2M</td><td>-0.59872</td><td>0.31021</td><td>-1.93</td><td>0.0536</td><td>.</td></tr>
<tr><td>Kur_ALLSKY_SFC_PAR_TOT</td><td>-0.2493</td><td>0.17578</td><td>-1.418</td><td>0.1561</td><td></td></tr>
<tr><td>Kur_PS</td><td>-0.36779</td><td>0.37617</td><td>-0.978</td><td>0.3282</td><td></td></tr>
<tr><td>severity_categoryLow</td><td>-0.12791</td><td>0.3878</td><td>-0.33</td><td>0.7415</td><td></td></tr>
<tr><td>Commercial.small.scaleCommercial</td><td>-0.12935</td><td>0.3333</td><td>-0.388</td><td>0.6979</td><td></td></tr>
<tr><td>TimeMid</td><td>-0.75632</td><td>0.33563</td><td>-2.253</td><td>0.0242</td><td>*</td></tr>
<tr><td>TimeFall</td><td>-0.71262</td><td>1.77766</td><td>-0.401</td><td>0.6885</td><td></td></tr>
<tr><th colspan="6">S4B Table. Coefficients Estimates and p-values SC2</th></tr>
<tr><td></td><td>Estimate</td><td>Std. Error</td><td>z value</td><td>Pr(>|z|)</td><td>Remarks</td></tr>
<tr><td>(Intercept)</td><td>5.649159</td><td>9.96304</td><td>0.567</td><td>0.57071</td><td></td></tr>
<tr><td>Rel</td><td>2.778071</td><td>1.414603</td><td>1.964</td><td>0.04955</td><td>*</td></tr>
<tr><td>Abs</td><td>-3.371384</td><td>2.463702</td><td>-1.368</td><td>0.17118</td><td></td></tr>
</table>

| | | | | | |
|---|---|---|---|---|---|
| Av_CLRSKY_SFC_PAR_TOT | -0.002903 | 0.067991 | -0.043 | 0.96594 | |
| Sd_T2M_RANGE | -0.632394 | 0.505208 | -1.252 | 0.21066 | |
| Sd_QV2M | -1.523832 | 0.488434 | -3.12 | 0.00181 | ** |
| Sd_RH2M | 0.332164 | 0.18487 | 1.797 | 0.07238 | . |
| Skew_T2M | -0.468924 | 0.52509 | -0.893 | 0.37184 | |
| Skew_WS10M | -0.036748 | 0.428786 | -0.086 | 0.9317 | |
| Skew_WD10M | 0.381384 | 0.764005 | 0.499 | 0.61764 | |
| Kur_RH2M | -0.676742 | 0.308747 | -2.192 | 0.02839 | * |
| Kur_ALLSKY_SFC_PAR_TOT | -0.295783 | 0.178039 | -1.661 | 0.09664 | . |
| Kur_PS | -0.327899 | 0.355548 | -0.922 | 0.35641 | |
| severity_categoryLow | -0.258732 | 0.386183 | -0.67 | 0.50288 | |
| Commercial.small.scaleCommercial | 0.013054 | 0.327643 | 0.04 | 0.96822 | |
| TimeMid | -1.066536 | 0.349663 | -3.05 | 0.00229 | ** |
| TimeFall | -0.087171 | 1.755737 | -0.05 | 0.9604 | |
| **S4C Table. Coefficients Estimates and p-values for SC3** | | | | | |
| | Estimate | Std. Error | z value | Pr(>\|z\|) | Remarks |
| (Intercept) | 19.732032 | 9.956541 | 1.982 | 0.0475 | * |
| Rel | 8.844749 | 1.566941 | 5.645 | 1.66E-08 | *** |
| Abs | -8.57641 | 2.414963 | -3.551 | 0.000383 | *** |
| Av_CLRSKY_SFC_PAR_TOT | -0.050602 | 0.064893 | -0.78 | 0.435522 | |
| Sd_T2M_RANGE | -1.412742 | 0.603792 | -2.34 | 0.019295 | * |
| Sd_QV2M | -3.930121 | 0.446249 | -8.807 | < 2e-16 | *** |
| Sd_RH2M | 1.196302 | 0.16269 | 7.353 | 1.93E-13 | *** |
| Skew_T2M | -2.241685 | 0.556118 | -4.031 | 5.56E-05 | *** |
| Skew_WS10M | -1.090465 | 0.53729 | -2.03 | 0.042401 | * |
| Skew_WD10M | 1.224661 | 0.807517 | 1.517 | 0.129374 | |
| Kur_RH2M | -1.476635 | 0.302377 | -4.883 | 1.04E-06 | *** |
| Kur_ALLSKY_SFC_PAR_TOT | -1.047583 | 0.293319 | -3.571 | 0.000355 | *** |

| | | | | | |
|---|---|---|---|---|---|
| Kur_PS | -0.823154 | 0.470234 | -1.751 | 0.080028 | . |
| severity_categoryLow | 0.001182 | 0.453269 | 0.003 | 0.997919 | |
| Commercial.small.scaleCommercial | -0.341213 | 0.444322 | -0.768 | 0.442523 | |
| TimeMid | -3.312953 | 0.37149 | -8.918 | < 2e-16 | *** |
| TimeFall | -0.348062 | 1.522631 | -0.229 | 0.819186 | |
| **S4D Table. Coefficients Estimates and p-values for SC4** | | | | | |
| | Estimate | Std. Error | z value | Pr(>\|z\|) | Remarks |
| (Intercept) | -0.80404 | 14.71984 | -0.055 | 0.956439 | |
| Rel | 1.24151 | 1.4429 | 0.86 | 0.389554 | |
| Abs | -3.07957 | 3.29072 | -0.936 | 0.349359 | |
| Av_CLRSKY_SFC_PAR_TOT | 0.04533 | 0.09875 | 0.459 | 0.646182 | |
| Sd_T2M_RANGE | -0.72327 | 0.64783 | -1.116 | 0.264233 | |
| Sd_QV2M | -2.37482 | 0.69654 | -3.409 | 0.000651 | *** |
| Sd_RH2M | 0.83273 | 0.20357 | 4.091 | 4.30E-05 | *** |
| Skew_T2M | 1.39475 | 0.62647 | 2.226 | 0.025989 | * |
| Skew_WS10M | -0.32152 | 0.54231 | -0.593 | 0.553276 | |
| Skew_WD10M | -1.11592 | 1.03569 | -1.077 | 0.281272 | |
| Kur_RH2M | -2.45601 | 0.4492 | -5.468 | 4.56E-08 | *** |
| Kur_ALLSKY_SFC_PAR_TOT | -0.24931 | 0.19982 | -1.248 | 0.212158 | |
| Kur_PS | 0.87233 | 0.51984 | 1.678 | 0.093332 | . |
| severity_categoryLow | 0.39417 | 0.62582 | 0.63 | 0.528795 | |
| Commercial.small.scaleCommercial | 2.04878 | 0.5189 | 3.948 | 7.87E-05 | *** |
| TimeMid | -1.8571 | 0.39085 | -4.751 | 2.02E-06 | *** |
| TimeFall | 0.78756 | 2.26932 | 0.347 | 0.728557 | |
| **S4E Table. Coefficients Estimates and p-values for SC5** | | | | | |
| | Estimate | Std. Error | z value | Pr(>\|z\|) | Remarks |
| (Intercept) | 11.21706 | 10.16496 | 1.104 | 0.2698 | |

| | | | | | |
|---|---|---|---|---|---|
| Rel | 2.50691 | 1.55908 | 1.608 | 0.1078 | |
| Abs | -3.37593 | 2.81359 | -1.2 | 0.2302 | |
| Av_CLRSKY_SFC_PAR_TOT | -0.06693 | 0.0697 | -0.96 | 0.3369 | |
| Sd_T2M_RANGE | -0.15072 | 0.56439 | -0.267 | 0.7894 | |
| Sd_QV2M | -0.64704 | 0.48031 | -1.347 | 0.1779 | |
| Sd_RH2M | 0.18163 | 0.17735 | 1.024 | 0.3058 | |
| Skew_T2M | -0.70121 | 0.57029 | -1.23 | 0.2189 | |
| Skew_WS10M | 0.09172 | 0.42188 | 0.217 | 0.8279 | |
| Skew_WD10M | 1.0561 | 0.81053 | 1.303 | 0.1926 | |
| Kur_RH2M | -0.36492 | 0.34108 | -1.07 | 0.2847 | |
| Kur_ALLSKY_SFC_PAR_TOT | -0.1519 | 0.19461 | -0.781 | 0.4351 | |
| Kur_PS | -0.42237 | 0.37508 | -1.126 | 0.2601 | |
| severity_categoryLow | -0.07415 | 0.39524 | -0.188 | 0.8512 | |
| Commercial.small.scaleCommercial | 0.13648 | 0.35294 | 0.387 | 0.699 | |
| TimeMid | -0.72787 | 0.34634 | -2.102 | 0.0356 | * |
| TimeFall | -1.8886 | 1.8353 | -1.029 | 0.3035 | |
| **S4F Table. Coefficients Estimates and p-values for SC6** | | | | | |
| | Estimate | Std. Error | z value | Pr(>\|z\|) | Remarks |
| (Intercept) | 18.39527 | 11.45125 | 1.606 | 0.1082 | |
| Rel | 3.46485 | 1.56161 | 2.219 | 0.0265 | * |
| Abs | -6.44961 | 2.95595 | -2.182 | 0.0291 | * |
| Av_CLRSKY_SFC_PAR_TOT | -0.09554 | 0.07417 | -1.288 | 0.1977 | |
| Sd_T2M_RANGE | 0.25179 | 0.57534 | 0.438 | 0.6617 | |
| Sd_QV2M | -0.84794 | 0.52218 | -1.624 | 0.1044 | |
| Sd_RH2M | 0.03155 | 0.17625 | 0.179 | 0.8579 | |
| Skew_T2M | -0.15048 | 0.5738 | -0.262 | 0.7931 | |
| Skew_WS10M | 0.9596 | 0.49637 | 1.933 | 0.0532 | . |
| Skew_WD10M | 1.40176 | 0.70708 | 1.982 | 0.0474 | * |

| | | | | | |
|---|---|---|---|---|---|
| Kur_RH2M | -0.02858 | 0.35188 | -0.081 | 0.9353 | |
| Kur_ALLSKY_SFC_PAR_TOT | -0.44131 | 0.20349 | -2.169 | 0.0301 | * |
| Kur_PS | -0.75161 | 0.43299 | -1.736 | 0.0826 | . |
| severity_categoryLow | -0.2211 | 0.38453 | -0.575 | 0.5653 | |
| Commercial.small.scaleCommercial | -0.79084 | 0.45786 | -1.727 | 0.0841 | . |
| TimeMid | -0.52945 | 0.32266 | -1.641 | 0.1008 | |
| TimeFall | -3.49091 | 2.03182 | -1.718 | 0.0858 | . |
| **S4G Table. Coefficients Estimates and p-values for SC7** | | | | | |
| Variable | Estimate | Std. Error | z value | Pr(>\|z\|) | Remarks |
| (Intercept) | 7.24389 | 10.4145 | 0.696 | 0.4867 | |
| Rel | 2.690481 | 1.52982 | 1.759 | 0.0786 | . |
| Abs | -4.204125 | 2.729126 | -1.54 | 0.1234 | |
| Av_CLRSKY_SFC_PAR_TOT | -0.034579 | 0.070498 | -0.49 | 0.6238 | |
| Sd_T2M_RANGE | -0.183299 | 0.500654 | -0.366 | 0.7143 | |
| Sd_QV2M | -0.903015 | 0.473942 | -1.905 | 0.0567 | . |
| Sd_RH2M | 0.226684 | 0.166676 | 1.36 | 0.1738 | |
| Skew_T2M | -0.450058 | 0.559788 | -0.804 | 0.4214 | |
| Skew_WS10M | 0.067903 | 0.415578 | 0.163 | 0.8702 | |
| Skew_WD10M | 0.553132 | 0.710031 | 0.779 | 0.436 | |
| Kur_RH2M | -0.495875 | 0.317551 | -1.562 | 0.1184 | |
| Kur_ALLSKY_SFC_PAR_TOT | -0.231255 | 0.183044 | -1.263 | 0.2065 | |
| Kur_PS | -0.242906 | 0.374147 | -0.649 | 0.5162 | |
| severity_categoryLow | 0.001393 | 0.382145 | 0.004 | 0.9971 | |
| Commercial.small.scaleCommercial | -0.132503 | 0.33316 | -0.398 | 0.6908 | |
| TimeMid | -0.782702 | 0.343206 | -2.281 | 0.0226 | * |
| TimeFall | -1.042053 | 1.846238 | -0.564 | 0.5725 | |

| S4H Table. Coefficients Estimates and p-values for SC8 | | | | | |
|---|---|---|---|---|---|
| Variable | Estimate | Std. Error | z value | Pr(>\|z\|) | Remarks |
| (Intercept) | 3.998924 | 10.20818 | 0.392 | 0.6953 | |
| Rel | 2.524384 | 1.477351 | 1.709 | 0.0875 | . |
| Abs | -3.500965 | 2.584967 | -1.354 | 0.1756 | |
| Av_CLRSKY_SFC_PAR_TOT | -0.007213 | 0.068945 | -0.105 | 0.9167 | |
| Sd_T2M_RANGE | -0.767154 | 0.511878 | -1.499 | 0.134 | |
| Sd_QV2M | -0.991548 | 0.466672 | -2.125 | 0.0336 | * |
| Sd_RH2M | 0.328385 | 0.16089 | 2.041 | 0.0412 | * |
| Skew_T2M | -0.958991 | 0.542482 | -1.768 | 0.0771 | . |
| Skew_WS10M | 0.273731 | 0.418942 | 0.653 | 0.5135 | |
| Skew_WD10M | 0.272908 | 0.697258 | 0.391 | 0.6955 | |
| Kur_RH2M | -0.582628 | 0.305984 | -1.904 | 0.0569 | . |
| Kur_ALLSKY_SFC_PAR_TOT | -0.214231 | 0.175512 | -1.221 | 0.2222 | |
| Kur_PS | -0.410215 | 0.380096 | -1.079 | 0.2805 | |
| severity_categoryLow | -0.208415 | 0.388656 | -0.536 | 0.5918 | |
| Commercial.small.scaleCommercial | 0.026641 | 0.333626 | 0.08 | 0.9364 | |
| TimeMid | -0.569569 | 0.333827 | -1.706 | 0.088 | . |
| TimeFall | -0.085233 | 1.76283 | -0.048 | 0.9614 | |

| S5 Table. Gene under positive selection pressure (Annotations according LH3 Genome) | | | | | | | | |
|---|---|---|---|---|---|---|---|---|
| **Pangenome_ta** | **Selection seasons** | **Description** | **Chromoson** | **start** | **end** | **strand** | **protein_id** | **locus_tag** |
| SUPERPANG_ ength=204874_ | 2020 Win 2021 Mid season | alpha/beta hydrolase | CP018475.1 | 90[illegible] | 90[illegible] | - | APO98621.1 | BJD13_0567 |

| | | | | | | | | |
|---|---|---|---|---|---|---|---|---|
| SUPERPANG_ ength=204874_ | 2022 Mid season,20 Winter | ClpV1 family T6SS ATPase | CP018475.1 | 96( | 96[illegible] | - | APO98662.1 | BJD13_0592 |
| SUPERPANG_ ength=204874_ | In all Seas | type VI secre system protei ImpG | CP018475.1 | 96[illegible] | 96[illegible] | - | APO98664.1 | BJD13_0593 |
| SUPERPANG_ ngth=1466113_ | In all Seas | S9 family peptidase | CP018475.1 | 270( | 27( | + | APP00022.1 | BJD13_1378 |
| SUPERPANG_ ength=281694_ | In all Seas | aminotransfer V | CP018475.1 | 307 | 30 | + | APP00309.1 | BJD13_1553 |
| SUPERPANG_ ength=281694_ | 2020 Win 2022 Win | fructose- bisphosphatas | CP018475.1 | 328( | 32[illegible] | + | APP00469.1 | BJD13_1648 |
| SUPERPANG_ ngth=1466113_ 5 | In all Seas | ClpV1 family T6SS ATPase | CP018475.1 | 371( | 37 | + | APP00778.1 | BJD13_1834 |
| SUPERPANG_ ngth=1466113_ 6 | In all Seas | S9 family peptidase | CP018475.1 | 383[illegible] | 38[illegible] | + | APP00863.1 | BJD13_1883 |
| SUPERPANG_ ngth=1466113_ | 2021 Mid season, 2( End-seaso | phosphometh rimidine synth ThiC | CP018475.1 | 449[illegible] | 45( | + | APP01419.1 | BJD13_2192 |
| SUPERPANG_ ength=76157_3 | In all Seas | TonB-depend receptor | CP018475.1 | 463( | 46[illegible] | + | APP01514.1 | BJD13_2247 |

| | | | | | | | | | |
|---|---|---|---|---|---|---|---|---|---|
| SUPERPANG_ ength=65201_1 | 2020 End season, 20 Winter | hypothetical protein | CP018472.1 | 7 | 7 | + | | APO97677.1 | BJD13_0034 |
| SUPERPANG_51_length=116847_81 | In all Seasons | type VI secretion system contractile sheath large subunit | Not found | | | | | | |
| SUPERPANG_51_length=116847_82 | In all Seasons | hypothetical protein XVE_0815 | Not found | | | | | | |
